# DICE: Fast and Accurate Distance-Based Reconstruction of Single-Cell Copy Number Phylogenies

**DOI:** 10.1101/2024.06.03.597037

**Authors:** Samson Weiner, Mukul S. Bansal

## Abstract

Somatic copy number alterations (sCNAs) are valuable phylogenetic markers for inferring evolutionary relationships among tumor cell subpopulations. Advances in single-cell DNA sequencing technologies are making it possible to obtain such sCNAs datasets at ever-larger scales. However, existing methods for reconstructing phylogenies from sCNAs are often too slow for large datasets. Moreover, the accuracies of many existing methods are highly sensitive to error and other features of the analyzed datasets.

In this work, we propose two new distance-based approaches for reconstructing single-cell tumor phylogenies from sCNA data. The new methods, *DICE-bar* and *DICE-star*, are based on novel, easy-to-compute distance measures and drastically outperform the current state-of-the-art in terms of both accuracy and scalability. Using carefully simulated datasets, we find that DICE-bar and DICE-star significantly improve upon the accuracies of existing methods across a wide range of experimental conditions and error rates while simultaneously being orders of magnitude faster. Our experimental analysis also reveals how noise/error in copy number inference, as expected for real datasets, can drastically impact the accuracies of many existing methods. We apply DICE-star, the most accurate method on error-prone datasets, to two real single-cell breast cancer datasets and find that it helps identify previously unreported rare cell populations.

## Introduction

Cancer progression is an evolutionary process driven by the accumulation of somatic mutations (Nowell 1976). Within tumors, there exist divergent subpopulations of cells characterized by distinct sets of somatic mutations, a phenomena in cancer called Intra-Tumor Heterogeneity (ITH) (Lawson et al. 2018). ITH is a primary obstacle in cancer prognosis, treatment, and prevention, and a better understanding of ITH is thought to be crucial for clinical success (Turajlic et al. 2019). One approach to studying ITH is through elucidating the evolutionary relationships between different cells in the tumor. Typically, the evolutionary history of a tumor is described by a *cell lineage tree*, where the leaves of the tree represent observed cells in the sample, and internal nodes represent ancestral cells (Beerenwinkel et al. 2014, Schwartz and Schäffer 2017). Rapid advances in high-throughput next-generation sequencing technologies have made it possible to infer such phylogenies (i.e., cell lineage trees) from a tumor’s mutational landscape. In particular, the recent development of single-cell DNA-sequencing (scDNA-seq) technologies has enabled the identification of individual cancer cell mutations at increasing scale and resolution. Such single-cell resolution data, despite technological limitations such as high error and dropout rates, shows great promise for inferring cell lineage trees and understanding ITH.

Somatic copy number alterations (sCNAs or simply CNAs) are the largest source of genetic heterogeneity in cancer genomes (Zack et al. 2013, Bignell et al. 2010, Beroukhim et al. 2010), making them valuable phylogenetic markers for reconstructing evolutionary trees. However, the properties of CNAs make tree inference challenging; for instance, there is a strong statistical dependence between adjacent genomic loci, and multiple events can overlap the same genomic region. Moreover, there is poor understanding of the distributions that govern CNA rates, sizes, and types (Beerenwinkel et al. 2014, Navin 2014). Several pioneering studies that leveraged single-cell CNA profiles to build tumor cell lineage trees used traditional correlation distances, such as Euclidean, combined with standard phylogeny inference algorithms (Navin et al. 2011, Wang et al. 2014, Schwartz and Schäffer 2017, Minussi et al. 2021). Such approaches, however, are thought to be ill-suited for copy number profiles because they make a number of simplifying assumptions and lack a more nuanced probabilistic model of evolutionary distance (Beerenwinkel et al. 2014, Schwartz and Schäffer 2017), although no systematic evaluation has been performed.

More recently, a number of studies have approached this problem using a framework that explicitly models copy number evolution based on a minimum evolution criterion. These frameworks typically involve finding the Minimum Event Distance (MED), defined as the minimum number of segmental amplifications or deletions needed to transform one genome into another. The MEDICC algorithm was the first method that uses the MED to both phase allele-specific copy numbers and reconstruct a phylogenetic tree from the copy number profiles (Schwarz et al. 2014). Since then, the MED model has been the focus of numerous other studies. The problem of finding the minimum number of events needed to transform one copy number profile into another, with certain restrictions, was shown to be solvable in linear time (Zeira et al. 2017), but the original problem approached by MEDICC, which aims to find a tree that minimizes the MED distance globally, was shown to be NP-hard (El-Kebir et al. 2017). There have also been some recent generalizations of the MED model. In the work of Cordonnier and Lafond (2020), individual events can alter copy numbers by any amount and are assigned a positive cost, with the resulting problem being to find a minimum-cost sequence. Zeira and Raphael (2020) proposed weighting events based on their length, event, and type. The MEDALT algorithm (Wang et al. 2021) uses the MED to infer an aneuploidy lineage tree which describes the sequence of events required to evolve one genome into the next, and uses a statistical test to identify CNAs associated with lineage expansion. The MEDICC2 algorithm (Kaufmann et al. 2022) builds upon its predecessor, MEDICC, by explicitly modeling Whole-Genome Duplication (WGD) events and implementing strategies to improve performance. Finally, the Lazac algorithm (Schmidt et al. 2023) solves a relaxation of the small parsimony problem under an approximation of the MED model that allows for the amplification of zero-copy regions.

One of the primary challenges in modeling copy number evolution under the MED framework is its underlying computational complexity. Current MED-based methods do not scale to large datasets, such as those that can be generated by modern scDNA-seq technologies, with many thousands of cells. A few approaches have sidestepped this concern by considering only breakpoints, the genomic locations joining adjacent segments with differing copy number. This idea was first developed in the context of bulk-sequencing data to compute clonal relationships between tumor samples of the same patient (Letouzé et al. 2010). Breakpoints have also been used in the Bayesian inference procedure of sitka (Salehi et al. 2023), as well as in an efficient approximation algorithm for the MED problem (Cordonnier and Lafond 2020). Overall, despite their challenges with scalability, MED-based approaches are currently the gold standard for reconstructing tumor cell lineage trees from single-cell CNA data.

In this work, we introduce two new methods, *DICE-bar* and *DICE-star*, based on novel, easy-to-compute distance measures, that drastically outperform the current state-of-the-art in terms of both accuracy and scalability. DICE-bar (short for “Distance-based Inference of Copy-number Evolution using breakpoint-root distance”) is a “CNA aware” approach that utilizes breakpoints between adjacent copy number bins to estimate the number of CNA events. In contrast, DICE-star (short for “Distance-based Inference of Copy-number Evolution using standard-root distance”) utilizes a simple penalized Manhattan distance between the copy number profiles themselves. Both methods then use the well-established balanced Minimum Evolution criterion (Desper and Gascuel 2002) to reconstruct the final tumor cell lineage tree. Remarkably, using a large number of realistically simulated datasets, we find that both DICE-bar and DICE-star consistently outperform the other methods across a wide range of experimental conditions, including different scales, resolutions, noise models, and error rates, while being orders of magnitude faster and more scalable than MED-based methods. Specifically, we find that (i) DICE-bar improves upon the accuracies of all MED-based methods across nearly all tested experimental conditions on both noise-free and noisy data, and (ii) DICE-star further dramatically improves upon the accuracies of all existing methods (including DICE-bar), resulting in up to 40% reduction in reconstruction error, on datasets with noise/error levels similar to those observed in real copy number profiles. Crucially, these findings hold true even for datasets generated using older, simpler simulators utilized by existing MED-based methods in their own evaluations.

Our comprehensive experimental analysis identifies DICE-star, given its exceptional tolerance to noise, as the most accurate tumor cell lineage tree inference approach, by far, for real scDNA-seq-based datasets. Remarkably, DICE-star’s improvements in accuracy over MED-based and other competing methods become even more pronounced as more realistic rates are used for key simulation parameters. Our results also show that DICE-bar, while not as accurate as DICE-star on noisy/error-prone data, provides the best performance among all CNA-aware methods, showing much better accuracy than the far more complicated MED-based methods. These findings are both surprising and significant given the simplicity and scalability of DICE-bar and DICE-star, and since distance-based phylogenetic approaches have traditionally been thought to be ill-suited for copy number profiles because they do not account for the specific mechanisms of copy number evolution (Beerenwinkel et al. 2014). Our results also clearly demonstrate the drastic effect that noise in copy number profiles has on the ability of MED-based methods, based on nuanced models of copy number evolution, to effectively reconstruct the underlying phylogeny. To assess its impact in practice, we applied DICE-star to two real scDNA-seq breast cancer datasets previously studied by Navin et al. (2011). On both datasets, the cell lineage trees computed by DICE-star identify possible rare cell populations that were not previously reported. These findings highlight the potential real-world impact of the proposed methods. DICE, an umbrella program implementing DICE-star, DICE-bar, and several other distance-based variants, is freely available open-source from https://github.com/samsonweiner/DICE.

## Materials and Methods

### Basic definitions

We assume a reference genome consists of *K* chromosomes where each chromosome *k* is partitioned into *n_k_* ordered bins labelled 1*,…,n_k_*. Here, each bin label corresponds to a unique contiguous subsequence from the reference. Any sampled genomes are aligned to the reference such that all genomes have the same number and sizes of bins for each chromosome. We refer to the number of times a bin appears in a sampled genome as that bin’s copy number, the values of which can be estimated across all samples and bins using methods for detecting CNAs from scDNA-seq data (Mallory et al. 2020, Wang et al. 2020). The *copy number profile (CNP)* for the *k*th chromosome is a vector *C^k^* = (*c*_1_*,…,c_nk_*) of non-negative integers, where *c_i_* denotes the copy number of bin *i* from that chromosome. A whole genome *s* can be described by a set of CNPs 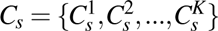. When considering whole human genomes without allosomes, *K* = 22 for total copy numbers and *K* = 44 for allele-specific copy numbers.

We define a *breakpoint* to be the difference between two consecutive copy numbers in a CNP *C*. More specifically, the breakpoint *b_i_* = *c_i_*_+1_ *−c_i_*, where 1 *≤ i ≤ n−* 1 and *n* is the number of bins in CNP *C*. A *breakpoint profile* is a vector *B^k^* = (*b*_1_*,…b_nk−_*_1_) obtained from a CNP *C^k^* that encodes the breakpoints of all consecutive bins from chromosome *k*.

### Description of DICE-bar and DICE-star

Consistent with other methods, we assume that the evolutionary history of a sampled population of cells can be described by a binary phylogenetic tree, where leaf nodes correspond to the observed single-cell genomes, and internal nodes represent the genomes of ancestral cells. This tree can be rooted along the branch leading to normal (non-cancer) cells or can be left unrooted.

Observe that cells that diverge later during tumor evolution are expected to have many shared alterations present in their genomes, while cells that diverge earlier will have comparatively fewer alterations in common. Thus, the evolutionary relationships among a set of sampled cells can be estimated by comparing the alterations present in their genomes. Given two genomes *s* and *t*, we can define a distance function *d* over their CNPs or breakpoint profiles, such that *d*(*s,t*) provides an estimate of the relative evolutionary distance between *s* and *t*. DICE-bar and DICE-star are both based on a simple, easy-to-compute distance functions to estimate relative evolutionary distances between sampled cells and use an off-the-shelf distance-based phylogeny reconstruction method to reconstruct the final tumor cell lineage tree based on the computed pairwise distances between sampled cells. Figure 1 provides an overview of DICE-bar and DICE-star and shows the key steps in their workflow. As the figure shows, both methods use the same novel distance function but DICE-bar applies this distance function to breakpoint profiles while DICE-star applies it directly to the input CNPs.

**Figure 1.**
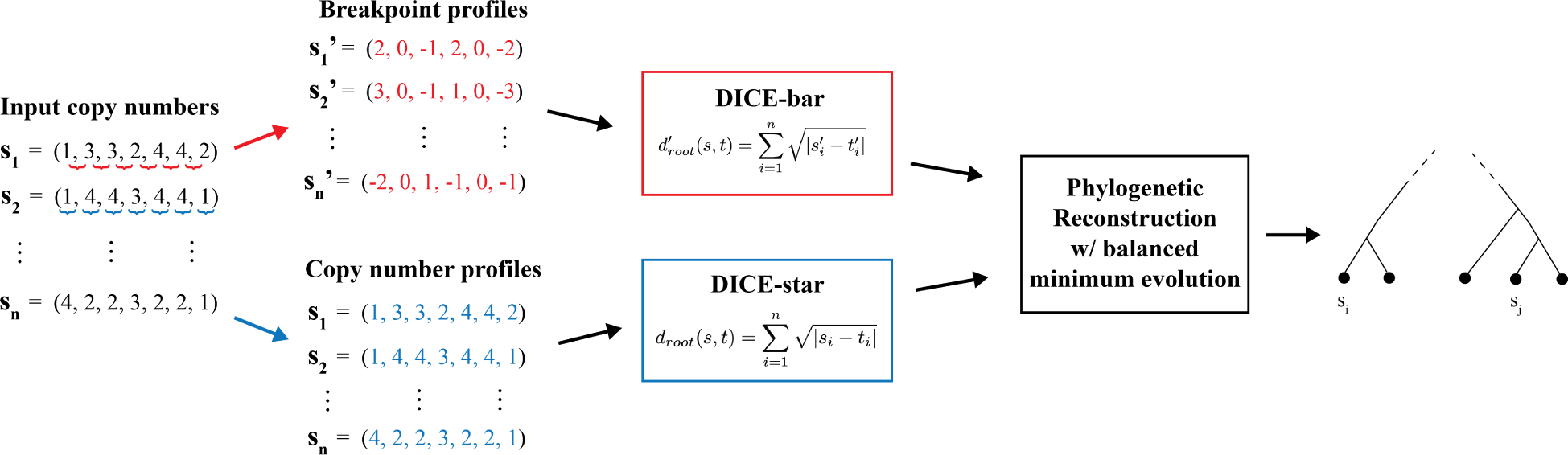
DICE-bar and DICE-star. DICE-bar and DICE-star both reconstruct tumor cell lineages from single-cell copy number profiles (CNPs) provided as input. Both methods employ the ‘root’ distance function as shown, with DICE-star applying it to the CNPs directly (standard variant; shown in blue), and DICE-bar applying it to the relative change in copy number at the breakpoints between adjacent genomic bins (breakpoint variant; shown in red). Once the pairwise distance matrix between cells has been computed, both DICE-star and DICE-bar use the ‘balanced minimum evolution’ criteria to reconstruct a cell-lineage tree.

#### DICE-star distance function

Given two genomes *s* and *t* with *K* chromosomes each, DICE-star uses the following distance function:

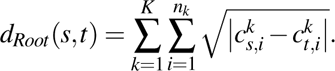

Note that this distance function is naive to the particularities of copy number evolution. Nonetheless, standard Euclidean and Manhattan distances between CNPs have been used previously for tree reconstruction of real tumor samples by Navin et al. (2011) and Minussi et al. (2021), respectively. Our novel ‘Root’ distance function above essentially applies the square root to each term of the Manhattan distance. This Root distance function is motivated by the following insight: CNAs can amplify a region by multiple copies in a single event, and thus larger changes in copy number do not necessarily imply greater evolutionary distances. Under the standard Manhattan distance, large changes are weighted equivalently to a number of events equal to its magnitude, potentially resulting in misleading evolutionary scenarios. At the same time, it is believed that the probability of a copy number amplification occurring scales inversely with its magnitude (Lauer et al. 2018, Liu et al. 2009). Accordingly, the Root distance attempts to better balance the effect of many low-magnitude CNA events versus few high-magnitude CNA events.

#### DICE-bar distance function

Observe that CNAs induce a strong statistical dependence among adjacent loci and multiple events may overlap. This suggests a number of pitfalls with using CNPs directly. First, the length of a CNA determines how many bins will be altered, essentially acting as a weight. Consider the scenario where two genomes differ by a single long event spanning many bins versus if they differed by many small events. It is more likely that the genomes are more closely related if they differ by a single event versus many, however if the long event exceeds the combined length of the small events, this will not be reflected in their computed distance. Second, independent events that overlap may be weighted less or nearly canceled out. This could occur if a region experiences a gain followed by a deletion event.

DICE-bar addresses this limitation by considering breakpoints rather than the copy numbers themselves. The use of breakpoints enables a more tailored metric for copy number evolution while still allowing for each site to be treated independently, and thus retaining the efficiency of a naive distance measure. In particular, DICE-bar uses the same novel Root distance function used by DICE-star but applies it to breakpoint profiles instead, as defined below:

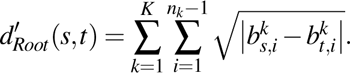

We note that the choice of distance function used by DICE-bar and DICE-star was based on a preliminary evaluation of several novel and existing alternatives; the results of that preliminary evaluation, based on a small subset of our simulated datasets, appear in Supplementary Section S.1.

#### Reconstructing final tumor cell lineage tree

Once all pairwise distances have been computed using the chosen distance function, the next step is to use a distance-based phylogenetic reconstruction approach to obtain the final cell-lineage tree. Both DICE-star and DICE-bar use balanced minimum evolution (Desper and Gascuel 2002), as implemented in the FastME software package (Lefort et al. 2015) to reconstruct the final cell-lineage tree. This choice of using balanced minimum evolution was based on a preliminary assessment, using a small subset of our simulated datasets, where we evaluated four possible distance-based phylogenetic reconstruction algorithms: Neighbor Joining (Saitou and Nei 1987), unweighted Neighbor Joining (Gascuel et al. 1997), balanced Minimum Evolution (Desper and Gascuel 2002), and ordinary least-squares Minimum Evolution (Rzhetsky and Nei 1993). Further details on the results of that assessment appear in Supplementary Section S.1.

### Experimental setup and evaluation metrics

We evaluate the performance of DICE-bar and DICE-star along with eight existing methods, MEDICC2 (Kaufmann et al. 2022), MEDALT (Wang et al. 2021), cnp2cnp (Cordonnier and Lafond 2020), Lazac (Schmidt et al. 2023), sitka (Salehi et al. 2023), WCND (Zeira and Raphael 2020), and the distance based approaches of Navin et al. (2011) and Minussi et al. (2021), using an extensive simulation study encompassing a wide variety of dataset types and conditions. The majority of our simulated datasets, consisting of simulated CNPs, were simulated using a recently published simulator called CNAsim (Weiner and Bansal 2023). CNAsim is the most sophisticated simulator currently available for simulating single-cell CNPs and implements a broad range of possible CNA mechanisms including whole genome duplication, whole-chromosomal CNAs, and chromosome-arm CNAs. CNAsim can also simulate clonal population structure through the accumulation of chromosomal CNAs and, crucially, implements a realistic error-model that (i) accounts for specific biases of single-cell sequencing that cause fluctuation in read counts, and (ii) models error patterns expected of existing CNA detection algorithms. For additional thoroughness, we also use datasets generated using simpler simulators developed and used by the authors of MEDICC2 (Kaufmann et al. 2022) and cnp2cnp (Cordonnier and Lafond 2020) to evaluate their own methods. We also apply the best-performing method for noisy datasets, DICE-star, to two real cancer datasets.

#### Overall experimental setup

We first evaluate DICE-bar, DICE-star, and the eight existing methods using CNAsim datasets simulated under wide range of parameter settings encompassing different experimental conditions, including different numbers of cells, numbers and lengths of chromosomes, CNA rates, copy number bin sizes, whole genome duplications, noise models, and error rates. Second, we use specially generated datasets to assess each method’s ability to accurately detect clones (sometimes also referred to as “subclones” in the literature), i.e., to group cells belonging to the same clone together on the reconstructed cell lineage tree. These special datasets were simulated to contain variable numbers of clones. Third, we evaluate DICE-star, DICE-bar, and the eight other methods on datasets simulated using two previous simulators developed and used in their own evaluation studies by the authors of MEDICC2 (Kaufmann et al. 2022) and cnp2cnp (Cordonnier and Lafond 2020). We use the same parameter settings used in the original studies to generate these additional datasets, enabling an “apples-to-apples” comparison of the methods, and, importantly, also explore the impact of using practicable ranges for key simulation parameters. Finally, we apply the best performing method, DICE-star, to two real publicly available breast cancer datasets from a previous study (Navin et al. 2011), analyze the resulting cell-lineage trees and contrast our findings against those reported previously.

#### Evaluation metrics

To evaluate the accuracies of different methods on the simulated datasets, we compare the cell lineage tree reconstructed by each method when applied to a simulated CNP against the known ground truth cell lineage tree used by the simulator for generating that CNP. Specifically, we use the well-known Robinson-Foulds (RF) distance (Robinson and Foulds 1981) between the reconstructed tree and the corresponding ground truth tree, assuming both trees to be unrooted. Briefly, the RF distance measures the number of bipartitions (or splits) that differ between the two phylogenetic trees being compared. Following standard practice, we normalize the RF distance to be between 0 and 1 by dividing the raw RF distance by the total number of non-trivial bipartitions in the two trees. Thus, a *Normalized RF distance (NRFD)* of 0 indicates the two trees are identical while an NRFD of 1 indicates that the two trees are maximally different (differing in all of their non-trivial bipartitions).

To assess the accuracy of clone detection, we measure how well a reconstructed cell-lineage tree identifies each clone (i.e., how well the cells belonging to that clone are grouped together in the reconstructed tree). Specifically, for any given ground truth clone, we identify the clade/bipartition in the reconstructed tree that shows maximum F1-score with respect to that ground truth clone. We then average these F1-scores across all clones present in the ground truth cell-lineage tree. Accordingly, a reported F1 score of 1 implies that all clones were detected with full accuracy.

#### Running existing methods

Of the eight existing methods included in our benchmark, MEDICC2, cnp2cnp, Lazac, and WCND all output binary cell lineage trees with observed cells as leaves. MEDICC2 is also capable of reconstructing ancestral copy number states, in addition to the cell lineage tree; this feature was disabled to enable a fair comparison of running times. For MEDALT and sitka, the output is a non-binary tree. The sitka tree outputs observed cells as leaves, so the evaluation metrics can be computed as-is. The MEDALT tree has observed cells as both internal and leaf nodes. To enable an apt comparison, we apply a transformation to the MEDALT tree that ensures each observed sample is represented as a leaf node. Finally, the methods of Navin et al. (2011) and (Minussi et al. 2021) do not have associated software implementations and we implemented them ourselves in the DICE software package. Further details on parameter settings and how each method was run appear in Supplemental Section S.4.

### Description of datasets

In the following, we first describe the extensive datasets simulated using CNAsim, then describe datasets simulated using the two older simulators, and finally describe the real cancer datasets used in this work.

### Datasets simulated using CNAsim

The majority of our datasets were simulated using the recently developed simulator CNAsim (Weiner and Bansal 2023), which simulates single-cell genomes along a ground truth cell lineage tree and can directly generate both noise-free and noisy CNPs. Next, we briefly describe the five key steps in the CNAsim simulation framework, discuss key simulation parameters, and justify our default values for these parameters.

1. *Generation of ground truth cell-lineage tree:* Following standard practice, a ground truth cell-lineage tree is simulated from an exponentially growing population under neutral coalescence (Hudson 2002, Posada 2020, Niida et al. 2020, Bozic et al. 2016, Beerenwinkel et al. 2007, Williams et al. 2016), the leaves of which correspond to observed cells in the experiment.
2. *Evolution of genome along cell-lineage tree:* An initial diploid genome is placed at the root of the tree and is represented by an ordered array of uniform-sized regions of *M* base-pairs each, where *M* is the minimum size of a CNA. Previous reports have classified variants as CNAs if its length exceeds 1000*bp* (Shlien and Malkin 2009, Feuk et al. 2006, Upadhyay et al. 2017, Pollex and Hegele 2007), which we use as a fixed value for *M*. The default number of chromosomes and chromosome lengths is derived from the GrHg38 human reference genome, however other custom genome sizes are considered. We add a number of focal CNAs to the root genome which represents a set of initial mutations that initiate cancer growth (by default, 10*×* that of normal edges; see below) (Sottoriva et al. 2015, Gao et al. 2016, Saito et al. 2018). Here, focal CNAs are defined as those smaller than a chromosome arm. Additionally, a whole-genome duplication (WGD) and/or chromosomal-CNAs can be introduced into the genome of the root cell (tumor founder) that are pervasive across the entire cell population (Bielski et al. 2018). Note that the only cell which can undergo WGD is the tumor founder cell. The genome is evolved along the tree topology, where each node inherits the genome of the parent node in addition to being altered by a Poisson-distributed number of focal CNAs. Existing estimates place the number of CNAs in human cancers to be in the many tens to hundreds (Gao et al. 2016, Minussi et al. 2021, Velazquez-Villarreal et al. 2020) for a single sample. Accordingly, we use a default of λ = 2 events per edge as this results in tens of focal CNAs globally for small trees and hundreds for large trees. Other values for this parameter are explored. Importantly, the total burden of events falls within ranges reported in pan-cancer studies (Harbers et al. 2021). A wide range of possible focal CNAs are generated by stochastically selecting for each event property. First, the paternal or maternal allele is selected with a draw from a Binomial distribution (default α = 0.5). The chromosome is selected at random with probability proportional to chromosome length. Each focal CNA is chosen to be either a copy number gain or deletion according to a draw from a Binomial distribution. Amplifications and deletions have been reported to occur in relatively equal numbers, albeit with high variance (Zack et al. 2013, Harbers et al. 2021). In general, these numbers tend to scale with one another (Beroukhim et al. 2010), and so we fix the probability of an amplification to be *p* = 0.5. The length of the CNA is drawn from an exponential distribution, as multiple studies have found the frequency of a CNA occurring scales inversely with its length (Itsara et al. 2010, Knouse et al. 2017). We fix the mean length of a CNA to be β = 5 Mbp based on existing estimates (Beroukhim et al. 2010, Gao et al. 2017, Harbers et al. 2021) while also allowing for meaningful evaluation. After a length is chosen, the starting location is selected on the chromosome uniformly among all possible locations. Lastly, if the event is a copy number gain, the number of additional copies is chosen with a draw from a geometric distribution (Mallory et al. 2020). The mean number of additional copies is fixed to be δ = 2 reflecting previous observations (Lauer et al. 2018, Liu et al. 2009), and all additional copies are inserted in tandem with the original.
3. *Larger-scale CNAs and inclusion of clonal structures:* Clonal structures can be added into the tumor population by selecting ancestral nodes whose clades represent diverging clones. In addition to being altered by focal CNAs, the genomes of selected ancestral nodes are subject to both whole-chromosomal and chromosome-arm CNAs, thereby significantly separating their lineage from the rest of the tree (Secrier et al. 2016, Zaccaria and Raphael 2021). To achieve this, a fixed number of ancestral nodes are chosen based on branch length and the size of the clade. To simulate chromosomal CNAs, we first determine if the event affects a chromosome arm or a whole chromosome with a Binomial distribution. We fix *p* = 0.75 with probability weighted in favor of chromosome-arm CNAs, as these are thought to be more common (Taylor et al. 2018). The specific chromosome or chromosome-arm is selected uniformly at random. Both chromosome-arm CNAs and whole-chromosomal CNAs can either be a deletion or duplication event, but not an amplification (Beroukhim et al. 2010), and is determined by a draw from a Binomial. By default, we set *p* = 0.5 for an even distribution of duplications and deletions. However, in the presence of WGD, we set *p* = 0.8 in favor of deletions, as the median ploidy of tumors having undergone WGD is close to 3 (Bielski et al. 2018). We note that clonal structures (or chromosomal CNAs) are not included in our default simulation parameter settings; instead we simulate additional datasets to explore the impact of varying numbers of clones.
4. *Generation of noise-free CNPs:* Upon completing the tree traversal, CNPs are generated for each leaf node by ‘mapping’ regions back to the starting diploid genome and grouping them into fixed-size bins. On real low coverage data, bin sizes typically are in the order of 500 kbp (Garvin et al. 2015) to 5 Mbp (Zaccaria and Raphael 2021). Accordingly, the default bin size is set to 1 Mbp, but we also evaluate other bin sizes.
5. *Generation of noisy CNPs:* In the last step, noise/error is introduced into the CNPs. This is important since copy numbers estimated from real sequencing data can be highly error-prone (Mallory et al. 2020). The simulator models two primary sources of noise in real CNPs. First, the ‘boundary’ model adds poor resolution at the edges of contiguous segments with the same copy number, which has previously been reported to be the main sources of error in copy number detection algorithms (Garvin et al. 2015, Mallory et al. 2020). Second, the ‘jitter’ model adds random fluctuations due to biases in sequencing technologies which often affect downstream analysis, for example uneven coverage (Navin 2014).

#### Selection of appropriate noise parameters

The benchmarking study of Mallory et al. (2020) evaluated the breakpoint detection accuracy of several existing CNA detection methods across a multitude of sequencing technologies and experimental conditions, and reported precision and recall values in the ranges of 0.4 *−* 0.75 and 0.5 *−* 0.8, respectively, for the best performing method. Based on this benchmarking study, we use two different noise levels, low and high, in our simulation. For the *low noise* setting, we used a boundary error rate, *r_b_*of 0.02 and a jitter error rate *r_j_* of 0.1, which results in a precision and recall of 0.710 and 0.761, respectively. For the *high noise* setting, we used *r_b_* = 0.04 and *r_j_* = 0.1, which results in precision and recall of 0.652 and 0.668, respectively. We note that while these parameter settings result in a significant loss in breakpoint detection accuracy, the vast majority of copy numbers remain unchanged because the number of breakpoints between contiguous segments is far less than the total number of bins. On average, only 0.064% of bins have an altered copy number in the low noise setting, and only 1.05% of bins have an altered copy number in the high noise setting. Further details on how these boundary and jitter error rates were selected and how precision and recall values were derived for the simulated noisy datasets appear in Supplementary Section S.2. In addition, we also generated datasets with varying jitter and boundary error rates to assess their impact on the reconstruction accuracies of different methods.

#### Simulation parameter ranges and defaults

To assess the robustness of the methods on different dataset types, scales, evolutionary conditions, error-rates, etc., we systematically explored the impact of changing key simulation parameter values. For each distinct combination of parameter settings, we generated 20 independent replicates and all reported results are averaged over these 20 replicates. A list of the key simulation parameters whose impact was systematically explored, along with their ranges and default values, appears below.

- Number of cells: *n ∈ {*10,25,50,100,250,500,1000,5000,10000*}*; default 250.
- Number of chromosomes: *x ∈ {*1,2,5,10,22*}*; default 22.
- Chromosome length (in Mbp): *y ∈ {*50, 100,200,500,750,1000*}*.

**–** If *x* = 22, then *y* uses lengths from the human reference genome hg38.
**–** If *x* =*̸* 22, then the default is *y* = 100 for every chromosome.
- Mean number of CNAs per edge: λ *∈ {*0.5,1,2,3,4,5*}*; default 2.
- Bin size (in kbp): *b ∈ {*500,1000,2000,5000,10000*}*; default 1000.
- Number of clones: *c ∈ {*0,2,4,6*}*; default 0.
- Whole-genome duplication: *w ∈ {*True, False*}*; default False.
- Boundary error rate: *r_b_∈ {*0,0.01,0.02,0.04,0.06,0.08,0.1,0.15*}*; default values of 0, 0.02, and 0.04 for “no noise”, “low noise”, and “high noise” datasets, respectively.
- Jitter error rate: *r_j_ ∈ {*0,0.05,0.1, 0.15*}*; default values of 0, 0.1, and 0.1 for “no noise”, “low noise”, and “high noise” datasets, respectively.

### Simulated datasets of existing methods

In addition to the datasets simulated using CNAsim, we also created simulated datasets using the simulation frameworks of the two existing methods MEDICC2 (Kaufmann et al. 2022) and cnp2cnp (Cordonnier and Lafond 2020). Both simulation frameworks take the same general approach as CNAsim; a ground truth tree topology is generated, CNAs are accumulated along each edge, and observed CNPs are derived from the genomes of the leaf nodes. However, there are several important differences between these two previous simulation frameworks and CNAsim related to number of bins, genome resolution, event sizes, event types, error models, etc. A description of the key differences appears in Supplementary Section S.3.

Using the simulation frameworks of MEDICC2 and cnp2cnp, we generated the same datasets as those outlined in their respective studies (Kaufmann et al. 2022, Cordonnier and Lafond 2020). We also used these older simulators to create additional datasets with more realistic parameter values for bin size and event rates.

### Real datasets

We applied our most robust method, DICE-star, to two publicly available cancer datasets. These datasets are derived from two breast cancer patients, T10 and T16, and consist of 100 cells each (Navin et al. 2011). The data for patients T10 and T16 were obtained as raw sequencing data using the SRA toolkit (Leinonen et al. 2010) in the form of fastq files for each cell. Following standard practices, sequencing reads were mapped to the hg38 human reference genome using the Burrows-Wheeler alignment tool (Li 2013), after which low-quality mapped reads were removed using Samtools (Li et al. 2009). The processed sequencing reads were sorted according to genomic position and passed as input to the SCOPE algorithm (Wang et al. 2020) to obtain the CNPs of each cell.

## Results

We present our results in six parts. First, we assess the accuracies of DICE-bar, DICE-star, and the eight existing methods using baseline simulated datasets, generated using default parameter settings and three different noise levels (no noise, low noise, high noise). For this assessment we evaluate both tree reconstruction accuracy and clone detection accuracy. Second, we perform a more in-depth evaluation of the methods by systematically assessing the impact of different experimental and evolutionary conditions, such as number of cells, number of CNA events, number of bins, etc., on their accuracies. Third, we more extensively assess the impact of CNA estimation error (i.e., noise) on the methods. Fourth, we evaluate the methods using the two previous simulators developed and used by the authors of MEDICC2 (Kaufmann et al. 2022) and cnp2cnp (Cordonnier and Lafond 2020). Fifth, we assess running times and scalability of the methods. And sixth, we present the results of applying DICE-star, the best performing method overall, to real breast cancer datasets.

### DICE-star and DICE-bar outperform other methods

We first used our baseline datasets, simulated using CNAsim with the default parameter values listed above and three different noise levels (no noise, low noise, and high noise), and additional datasets aimed at assessing the ability of the different methods to accurately detect clones, to assess all ten methods. The error-rates for low-noise and high-noise datasets were selected to match the breakpoint detection accuracy of existing CNA detection algorithms (Mallory et al. 2020), with low-noise datasets corresponding to the higher end of observed precision and recall values and high-noise datasets corresponding to the middle of observed ranges for precision and recall. Further details on error rates and noise parameters appear in Supplementary Section S.2 and Supplementary Table S1. No-noise datasets represent the ideal case when all copy number alterations are inferred without any error.

#### Cell lineage reconstruction accuracy

Figure 2 shows the accuracies of cell lineage reconstruction for all ten methods on the baseline datasets. For each of the three noise levels, reported accuracies are averaged over 20 datasets, where each dataset consists of 250 cells and uses 22 chromosomes with lengths based on the human reference genome hg38 (and default values for all other simulation parameters). As expected, we find that the presence of noise results in higher mean reconstruction error and overall variance for all methods. This analysis reveals important insights into the relative performance (accuracies) of the different methods. In particular, we find that (i) DICE-bar matches or improves upon the accuracies of all existing methods on the noise-free datasets, and significantly improves upon the accuracies of the more sophisticated methods MEDICC2, MEDALT, cnp2cnp, Lazac, and WCND even on the noisy datasets, and (ii) DICE-star shows exceptional robustness to noise and drastically improves upon the accuracies of all existing methods, as well as of DICE-bar, on all noisy datasets. For example, on the high-noise datasets, DICE-star and DICE-bar show normalized RF distances of 0.353 and 0.507, respectively, while the existing methods MEDICC2, MEDALT, cnp2cnp, Lazac, Sitka, WCND, Navin et al., and Minussi et al. show Normalized RF distances of 0.584, 0.861, 0.57, 0.763, 0.808, 0.588, 0.662, and 0.384, respectively. Remarkably, DICE-star improves upon the normalized RF distances of the top performing sophisticated methods MEDICC2 and cnp2cnp by an average of 43% and 39.3%, respectively, for the noisy datasets.

**Figure 2.**
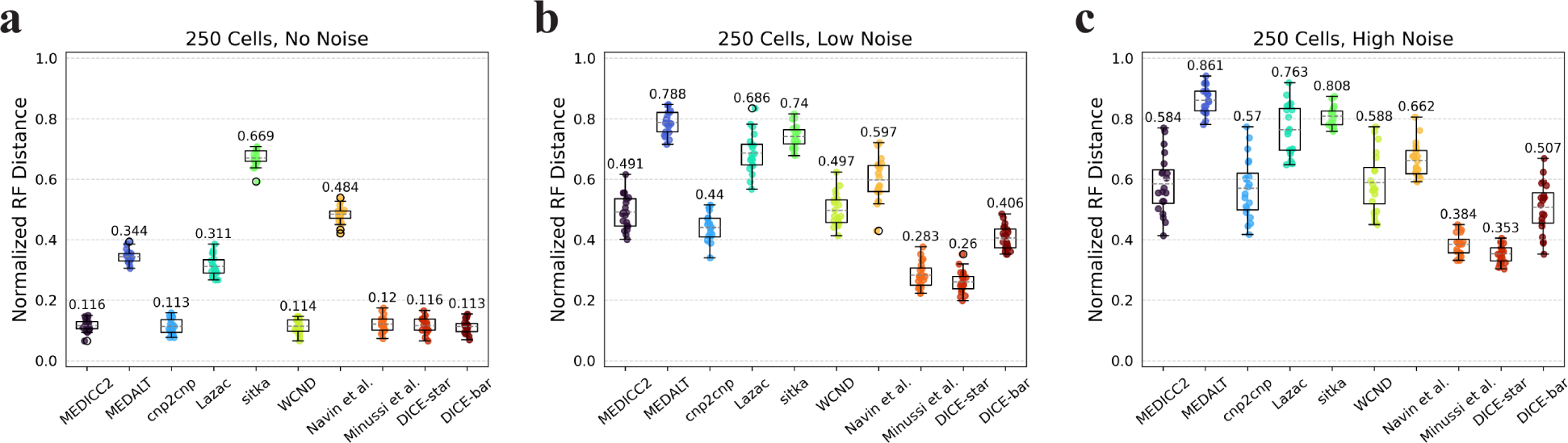
Cell lineage tree reconstruction accuracies. Box and whisker plots are shown for DICE-bar, DICE-star, and eight existing methods on simulated datasets with 250 cells and varying levels of noise. Results are shown for datasets with: (**a**) no noise; (**b**) low noise; (**c**) high noise. Lower Normalized RF Distances imply greater reconstruction accuracy. Data was generated using CNAsim and default simulation settings, and error rates for low and high noise levels were selected to match the precision and recall characteristics of breakpoint detection from existing CNA detection algorithms. Observe that DICE-bar matches or improves upon the accuracies of all other methods on noise-free datasets and DICE-star drastically improves upon the accuracies of all other methods on the noisy datasets.

More generally, these results suggest that methods based on MED or breakpoint distances, though accurate on noise-free data, can be highly sensitive to error and noise in inferred CNPs. Likewise, these results also show that methods based on appropriately designed distance measures between CNPs, such as DICE-star and the method of Minussi et al. (which uses standard Manhattan distance between CNPs), though not as accurate as the best MED or breakpoint methods on noise-free data, are far more robust to realistic levels of noise and error in CNPs. These findings have important implications for the application of these methods to real datasets.

#### Clone detection accuracy

To evaluate clone detection, we generated datasets containing a variable number of clones in the ground truth tree, with all other parameters kept at default values. Figure 3 shows the results of DICE-bar, DICE-star, and existing methods on trees containing exactly four clones. On noise-free data, we find that most methods perform exceptionally well at capturing clonal populations. In particular, both DICE-bar and DICE-star as well as MEDICC2 and the method of Minussi et al. obtain perfect or near-perfect scores. On noisy data, most methods show a decrease in performance, with DICE-bar and the MED-based methods MEDICC2, MEDALT, cnp2cnp, Lazac, and WCND all showing clear drops in clone detection accuracy. However, DICE-star and the method of Minussi et al. show exceptional robustness to noise, outperforming all other methods, including MEDICC2, at both low and high levels of noise. For example, both DICE-star and the method of Minussi et al. show an F1 score of 1.0 on the high-noise datasets, while MEDICC2, MEDALT, cnp2cnp, Lazac, sitka, WCND, Navin et al., and DICE-bar show F1 scores of 0.963, 0.63, 0.924, 0.851, 0.851, 0.93, 0.957, and 0.949, respectively. Sitka also appears to be robust to noise, likely due to its loci preprocessing step, but has an overall poor baseline performance compared to the other methods. Overall, these results further substantiate the superiority of DICE-star for real datasets.

**Figure 3.**
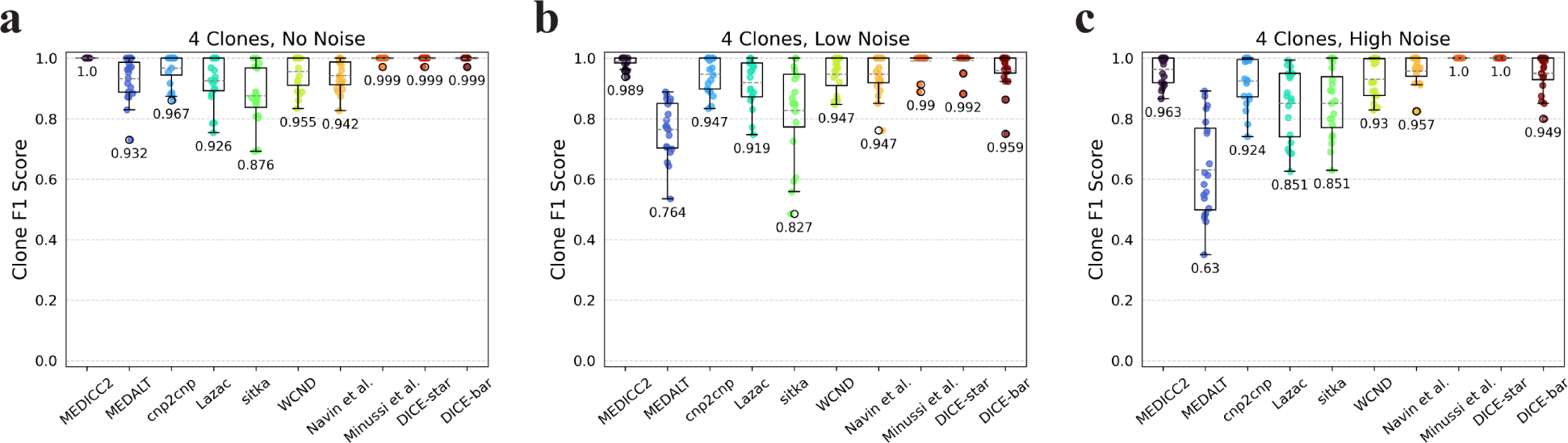
Clone detection accuracies. Box and whisker plots of F1 scores are shown for DICE-bar, DICE-star, and eight existing methods on simulated datasets with 250 cells and varying levels of noise. Scores are averaged over 20 datasets, each with exactly four clones. Results are shown for datasets with: (**a**) no noise; (**b**) low noise; (**c**) high noise. Higher F1 scores are better. As the plot shows, DICE-bar, DICE-star, MEDICC2, and Minussi et al. show perfect or near-perfect F1 scores on noise-free data, while DICE-star and Minussi et al. exceed the F1 scores of all other methods on the noisy datasets.

Importantly, we found that clone detection performance remains virtually unchanged when varying the number of clones (Supplementary Figure S3). We also note that the presence of clones has little to no effect on overall cell lineage reconstruction accuracy (Supplementary Figure S2).

### DICE-star and DICE-bar are robust to evolutionary and experimental conditions

Next, we evaluated the methods on datasets representing different evolutionary and experimental conditions such as different numbers of cells, genome/chromosome sizes, CNA rates, bin sizes, etc. For each condition, we varied the relevant simulation parameter, keeping other parameters at their default values, and generated no-noise and high-noise datasets.

#### Number of cells

We evaluated the impact of number of cells (i.e., tree size) using datasets with 10, 25, 50, 100, 250, and 500 cells. Supplementary Figures S4 and S5 show the results of this analysis. Consistent with previous results, we find that DICE-bar outperforms all other methods on all noise-free datasets except for the one with 25 cells where, unexpectedly, DICE-star becomes the best performing method. For the noisy datasets, we again find that DICE-star drastically outperforms all other methods, including DICE-bar. Overall, we find that the accuracy of most methods is only slightly affected by the number of cells, but that variance across replicates consistently decreases with increasing numbers of cells. The one exception to this is sitka, which achieves competitive performance on small trees only and, interestingly, shows robustness to noise, even slightly outperforming DICE-star in the 25 cell setting. This is likely due to the highly selective loci filtering step employed by sitka, which greatly restricts the number of loci used for phylogenetic reconstruction. The small number of filtered loci is only sufficient for reconstructing trees for small numbers of cells.

#### Genome size

We next evaluated the impact of genome size by generating datasets with varying number of chromosomes, each with a length of 100 Mbp, and keeping all other parameters at default values. Supplementary Figure S6 shows the results of this analysis. On the noise-free datasets, we find that all methods perform worse with fewer chromosomes and that, as expected, DICE-bar outperforms all other methods. On the noisy datasets, DICE-star significantly outperforms all other methods for all numbers of chromosomes. Surprisingly, we also find that the performance of all methods slightly worsens as the number of chromosomes increases from 5 to 10 for the noisy datasets. This may be an artifact of the error model used in CNAsim, or because the methods see no additional benefit from additional error-prone CNPs beyond 5 chromosomes given that the number of cells in these datasets is only 250. For completeness, we also performed an equivalent experiment where the number of chromosomes was fixed at 1, but chromosome length varied; for example, 5 chromosomes with length 100 Mbp would correspond to 1 chromosome with length 500 Mbp. We found nearly identical results across the two sets of experiments (Supplementary Figure S7), suggesting that the total amount of available genomic information is the more impactful factor, regardless of how it is distributed across chromosomes.

#### Presence of WGD

We find that the presence of WGD has little impact on reconstruction accuracy in the noise-free setting, but that all methods show worse performance with WGD (than without WGD) on the noisy datasets (Supplementary Figure S8). In addition, as the figure shows, most methods appear to improve slightly if the WGD is followed by chromosomal CNAs. Overall, DICE-bar and DICE-star remain the best performing methods, by far, on noise-free and noisy datasets, respectively. Interestingly, MEDICC2 becomes the best performing method if there is no noise and the WGD is introduced as the sole large-scale copy number event. This is not very surprising since MEDICC2 is the only method that makes an explicit attempt to model WGD. Still, DICE-star greatly outperforms MEDICC2 on all noisy datasets, cutting its reconstruction error almost in half.

#### Number of CNA events

We varied the mean number of focal CNAs per edge (parameter λ) in the range 0.5 to 5 and found it to be among the most impactful parameters for cell lineage reconstruction accuracy. As Supplementary Figures S9 and S10 show, increasing CNA event rates lead to significant improvements in reconstruction accuracies for all methods. On noise-free datasets, MEDICC2, cnp2cnp, WCND, Minussi et al., DICE-star, and DICE-bar all achieve near-perfect reconstruction accuracy when λ is set to 5 (Supplementary Figure S9). This is likely because the abundance of mutations along each edge substantially reduces the uncertainty of evolutionary relationships. Interestingly, on noise-free datasets, MEDICC2 outperforms all other methods for the two lowest values of λ, with DICE-bar showing best performance on all other noise-free datasets. This indicates that the MED model is best suited for low mutation rates. However, as expected, the performance of MEDICC2 falls sharply on noisy datasets, and DICE-star again shows the best performance across all CNA event rates on noisy datasets (Supplementary Figure S10).

#### Number of bins and bin size

Finally, we evaluated the impact of bin size (or, equivalently, number of bins) by considering four different bin sizes in the range 0.5 Mbp to 10 Mbp. Results are shown in Supplementary Figure S11. On the noise-free datasets, we find that accuracy steadily decreases with increasing bin size. This is expected since smaller bin sizes (which leads to a greater number of total bins) provide more information for phylogenetic reconstruction. However, we see mixed results on the noisy datasets, with only DICE-star and Minussi et al. clearly benefiting from smaller bin sizes. This suggests that the other methods, which are all more susceptible to noise, may be unable to benefit from smaller bin sizes on real, error-prone datasets. DICE-star remains the best method on noisy datasets across all bin sizes, and DICE-bar outperforms the existing methods on noise-free datasets for 3 of the 4 bin-sizes. On the noise-free dataset with largest bin-size, MEDICC2 shows slightly better accuracy than DICE-bar.

Overall, the above results demonstrate the robustness of DICE-star and DICE-bar to different evolutionary and experimental conditions and further establish DICE-bar as the best method to use for noise-free data and DICE-star as the best method to use for noisy data. These results also suggest that MEDICC2 does well in scenarios where there is limited information in the CNPs, e.g., when the rate of events is low or when bin sizes are very large, as long as the CNPs are noise-free.

### DICE-star is robust to CNA estimation error

To better understand the effect of CNA estimation error (i.e., noise in input CNPs) on cell lineage reconstruction accuracy, we generated datasets using several additional combinations of the noise parameters. Specifically, we fixed the jitter error rate at two values, 0.05 and 0.15, representing a low and high baseline rate, and varied the boundary error rate from a low of 0.02 to a high of 0.14 at intervals of 0.02. For reference, Supplementary Table S1 shows how these error rates affect the precision and recall of ground truth CNPs.

Figure 4 shows the results of applying all ten methods to these datasets. As the figure show, the performance of all methods degrades rapidly as the boundary error-rate increases. More importantly, regardless of the source of noise (i.e. the boundary model or jitter model), DICE-star outperforms all other methods at all noise levels. This improvement over other methods is greatest when the rate of jitter error is high, but the magnitude of improvement decreases with increasing boundary error rate. Interestingly, while MEDICC2, MEDALT, cnp2cnp, Lazac, WCND, and DICE-bar are all substantially negatively impacted by increased jitter, this is not the case for DICE-star and the method of Minussi et al. (2021). We also note that DICE-bar continues to improve upon the accuracies of the more sophisticated methods MEDICC2, MEDALT, cnp2cnp, Lazac, sitka, and WCND across all error rates.

**Figure 4.**
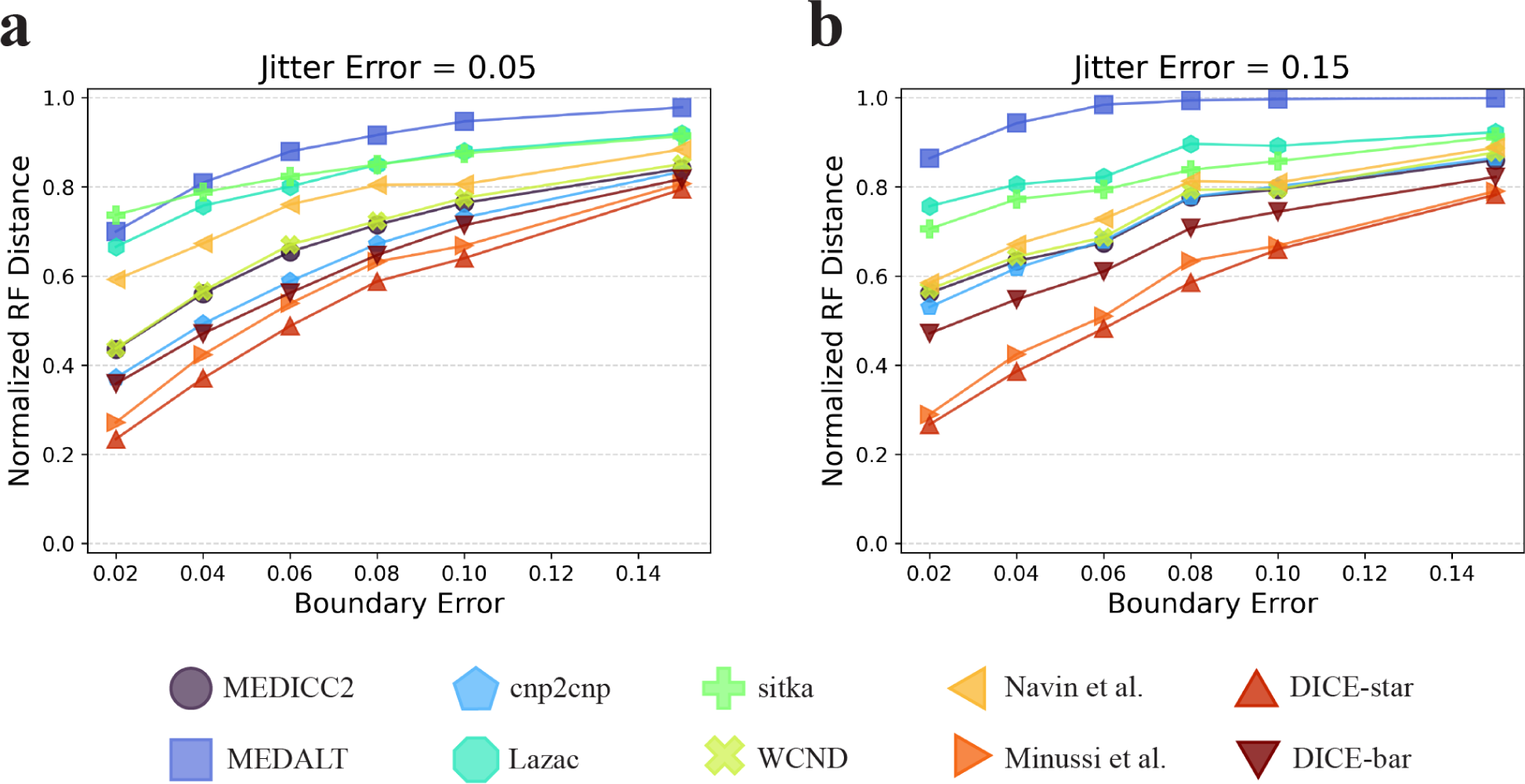
Impact of CNA estimation error on the different methods. The plots show how the individual error models (boundary and jitter) and their error rates affect cell lineage reconstruction accuracy. The parameter of the jitter model was fixed and various parameters were evaluated for the boundary model (*y*-axis). The jitter model was fixed at a ‘low’ error rate with *r_j_*= 0.05 (**a**) and a ‘high’ error rate with *r_j_* = 0.15 (**b**). For each jitter error rate the plots show reconstruction accuracy results for seven different boundary error rates. Observe that DICE-star outperforms all other methods for all error-rates, and that even DICE-bar outperforms the more sophisticated methods MEDICC2, MEDALT, cnp2cnp, Lazac, sitka, and WCND across all error-rates.

### Other simulators support exceptional performance of DICE-bar and DICE-star

To corroborate the above results and confirm that the relative accuracies of the different methods are not affected by the specifics of the simulation procedure used to generate CNPs, we generated additional simulated datasets following the simulation studies of existing methods. Specifically, we used the simulation frameworks of MEDICC2 (Kaufmann et al. 2022) and cnp2cnp (Cordonnier and Lafond 2020), both of which provide an implementation.

#### MEDICC2 simulator

We generated datasets of various sizes using the MEDICC2 simulator with its default parameters and with their most frequently used mutation rate of 0.05 (Kaufmann et al. 2022). These datasets consist of three sizes (100, 250, and 500 cells), and for each size includes datasets with no WGD and a high rate of WGD. We note that the MEDICC2 simulator cannot simulate noisy CNPs, and therefore all MEDICC2 datasets are noise-free. Supplementary Figure S13 shows the results of applying all ten methods to these datasets. Consistent with our previous results on noise-free datasets, DICE-bar shows substantially better accuracy than all other methods on both the no-WGD and high-WGD datasets. WCND and MEDICC2 also perform well on both the no-WGD and high-WGD datasets, while DICE-star and the method of Minussi et al. (2021) perform well on the no-WGD datasets. Interestingly, the presence of WGD appears to have a substantial negative impact on the performance of all methods, although MEDICC2, WCND, and DICE-bar appear to be less affected than other methods. We also note that sitka performs extremely poorly on these datasets, likely due to its strict loci filtering step. Overall, these results further confirm DICE-bar as the most accurate method on noise-free data.

For greater realism, we also used the MEDICC2 simulator to generate datasets with additional bins per chromosome. At 22 chromosomes, the MEDICC2 simulator by default only uses 10 bins per chromosome for a total of 2 *×* 22 *×* 10 = 440, which may not be realistic for whole-genome datasets. For reference, the default CNAsim parameters of using 1 Mbp bin sizes over the 22 autosome lengths derived from hg38 equates to over 5700 bins. Because the MEDICC2 simulator uses a number of mutations that scales linearly with the total number of bins, we used a scaled mutation rate to maintain consistency with default settings; however, for comparison we also show results using a fixed mutation rate. Results are shown in Supplementary Figure S14. We find that DICE-bar continues to be the best performing method, improving upon the accuracy of the nearest competitor, WCND, by at least 15%. Interestingly, while MEDICC2, cnp2cnp, sitka, and DICE-bar all show a steady improvement in performance as the number of bins increases with scaled mutation rates, the performance of the remaining methods remains relatively unchanged or slightly worsens. In contrast, genome size strongly correlated with performance across all methods in the CNAsim datasets (Supplementary Figures S6 and S7). When keeping the mutation rate fixed, which effectively increases the number of mutations, DICE-bar achieves near perfect reconstruction accuracy, however the other methods either show similar results to the scaled setting or become worse. This result is unexpected, as other methods do benefit from an increased mutation rate in the CNAsim datasets (Supplementary Figure S9 Parts (e), (f)). This discrepancy can likely be attributed to the many types of mutations outside of segmental duplications and deletions implemented in the MEDICC2 simulator.

#### cnp2cnp simulator

Next, we evaluated the different methods on 100-cell datasets generated with the cnp2cnp simulator using its default parameter values. This cnp2cnp simulator implements a simple error model to generate noisy CNPs, and Supplementary Figure S15 shows results on datasets with increasing levels of noise. As the figure shows, DICE-bar significantly outperforms all other methods on the two lowest noise datasets, while DICE-star begins to significantly outperform all other methods for the remaining three datasets with higher noise levels. These results are fully consistent with our previous findings. As before, DICE-bar also continues to outperform the more sophisticated methods MEDICC2, MEDALT, cnp2cnp, Lazac, sitka, and WCND across all noise levels. We also find that the performance of all methods is worse on the cnp2cnp datasets than on the baseline CNAsim datasets. This is likely due to the very low default number of bins, set to 100 on a single chromosome, in the cnp2cnp simulator.

We also used the cnp2cnp simulator to generate more realistic datasets with additional bins. Unlike the other two simulators, cnp2cnp uses genomes consisting of a single haploid chromosome, though there is a parameter controlling the number of bins (default 100). As Supplementary Figure S16 shows, the performance of all methods improves as the number of bins increases, with mean performances approaching those observed on the CNAsim datasets. Interestingly, at the two highest bin settings (1000 and 2000) and the lowest noise levels (0, 0.1), Lazac becomes the best performing method, very slightly outperforming DICE-bar. Otherwise, the relative performance of the methods remains mostly unchanged, with either DICE-bar or DICE-star outperforming the other methods depending on the level of noise in the dataset. We also find that, at the highest bin setting of 2000, the method of Minussi et al. (2021) matches the performance of DICE-star at all noise levels. This is likely an artifact of the mutation model of the cnp2cnp simulator being limited in how copy numbers change in individual events.

### DICE-bar and DICE-star are highly scalable

Besides demonstrating improved accuracy, DICE-bar and DICE-star have an exceptional advantage in scalability and computational efficiency, handling thousands of cells in a matter of minutes. Figure 5 reports running times of the different methods on noise-free datasets with varying numbers of cells (and generated using default values for other parameters). All methods were run on a single core of an Intel Xeon 2.1 GHz processor with 64 GB of RAM. As the figure shows, both DICE-bar and DICE-star, are much faster and more scalable than all MED-based methods. For example, DICE-bar and DICE-star are both over 1000 times faster than MEDICC2 on the 500-cell datasets. The methods of Navin et al. and Minussi et al., being similarly distance-based, have nearly identical running times as DICE-bar and DICE-star. Observe that sitka becomes the fastest method for 5000 cells; however, this result is misleading since the running time of sitka depends on the number of steps used in its Markov chain Monte Carlo search heuristic, which was kept constant across all runs.

**Figure 5.**
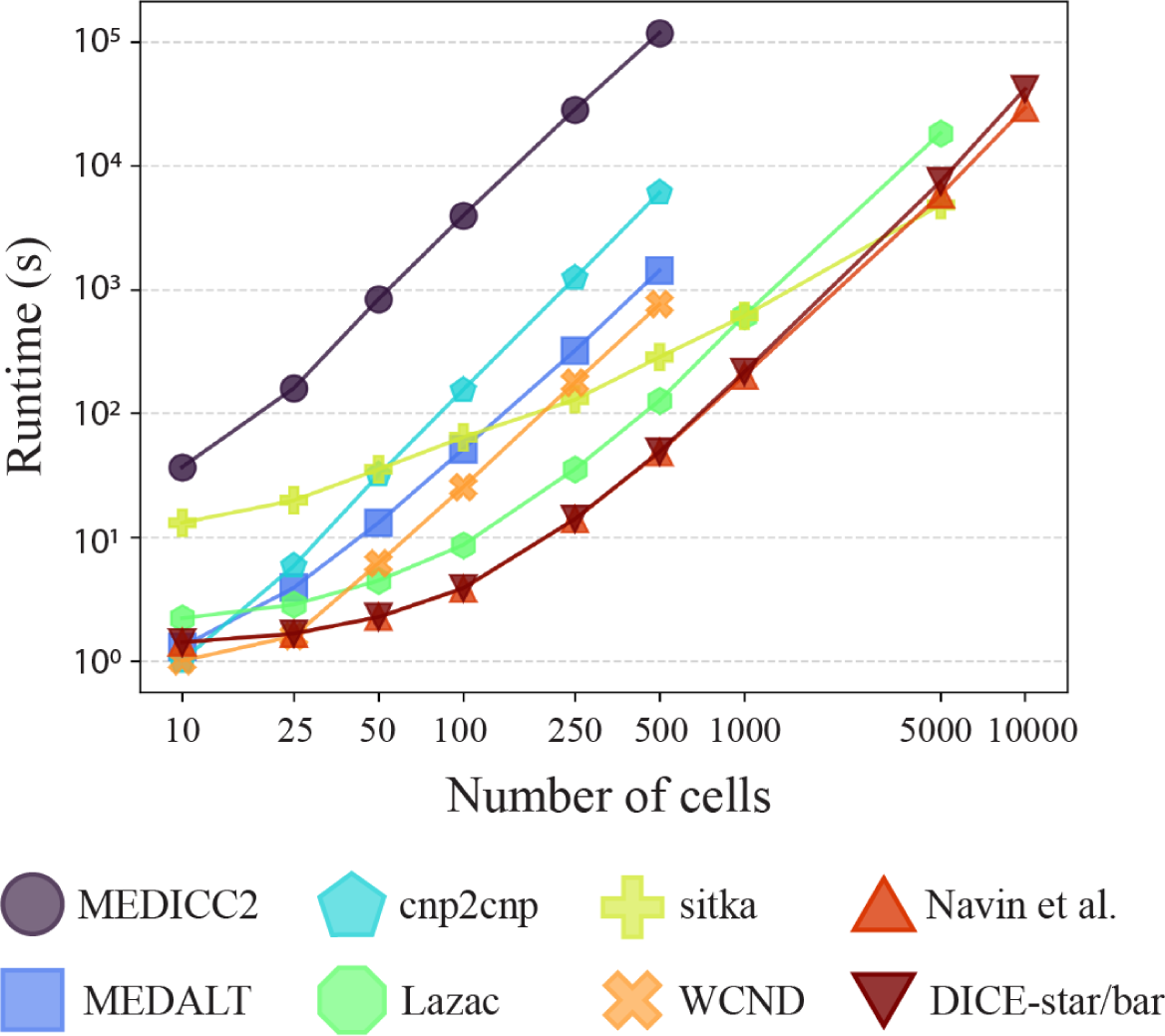
Running time and scalability. Running times are shown for MEDICC2, MEDALT, cnp2cnp, Lazac, sitka, WCND, Navin et al., Minussi et al., DICE-bar, and DICE-star on noise-free datasets with varying numbers of cells. Running times for DICE-bar, DICE-star, and the method of Minussi et al. (2021) are identical and are shown together. The *x*-axis is the log-scaled number of cells in the dataset, and the *y*-axis is the log-scaled runtime in seconds. Observe that DICE-star and DICE-bar are orders of magnitude faster and more scalable than the best MED-based methods. All reported times are averaged over 20 runs executed using a single core of an Intel Xeon 2.1 GHz processor.

We also assessed the impact of CNA estimation error (i.e., noise) on running times and found that all methods report a slight increase in running time (Supplementary Figure S12).

### DICE-star provides new insights into cell lineage evolution in two breast cancer datasets

Given the substantially superior accuracy of DICE-star on noisy data, we applied DICE-star to two previously published scDNA-seq datasets of high-grade triple negative ductal carcinomas, referred to as T10 and T16 (Navin et al. 2011). As we describe below, our analysis reveals new insights into the emergence and evolution of clonal populations in these datasets.

In the previous study of Navin et al. (2011), fluorescence-activated cell sorting (FACS) was used to study the distribution of ploidy across the single-cell populations, revealing a majority diploid fraction and smaller aneuploid fractions for both tumors, as well as a hypodiploid fraction in T10. Additionally, a small number of ‘pseudo-diploid’ cells were reported in both tumors, later characterized as having near-diploid aberrant CNPs with diverse gains and losses. Using this information, Navin et al. (2011) selected 100 flow-sorted single cells each from T10 and T16, taking care to include representatives from the various ploidy fractions and anatomical sectors. For T16, this included 48 cells from a paired metastatic liver carcinoma. To investigate population structure, Navin et al. (2011) constructed evolutionary trees using Neighbor Joining based on pairwise Euclidean distances between CNPs derived in situ (referred to as the method of Navin et al. in our simulation study), revealing large well-defined clades with relatively small inter-clade distances in both tumors. In particular, the subpopulations induced by the 4 major clades of their T10 tree map perfectly to the represented ploidy fractions, namely a diploid, hypodiploid, and two aneuploid populations. Similarly in their T16 tree, two major clades clearly separate the diploid and aneuploid populations, and the aneuploid clade is further subdivided into two smaller subpopulations, representing cells from the primary and metastatic sites. Importantly, all pseudo-diploid cells included in either of their trees appear in the diploid clade, and are seemingly unrelated to each other.

In our reanalysis of these datasets, we generated the CNPs by applying SCOPE (Wang et al. 2020) to the aligned and processed sequencing data using a bin size of 100 kbp. SCOPE, which employs a number of quality control and normalization features, removed 1 cell each from both tumors, and of the remaining 99 cells returned CNPs over 24779 and 24534 bins for T10 and T16, respectively. DICE-star cell lineage inference took mere seconds for both tumors on a personal laptop computer. Figure 6 and Supplementary Figure S17 show DICE-star trees for T10 and T16, respectively. Consistent with the cell lineage trees reported by Navin et al. (2011), the DICE-star trees of both T10 and T16 show divergent clades that strongly correspond to the same major clonal populations as Navin et al. (2011) and with the same macro-evolutionary relationships. However, unlike the cell lineage trees of Navin et al. (2011), both DICE-star trees contain additional small clades or single nodes that appear to diverge from the major clonal clades. These clades/single nodes, while having ploidies similar to that of the existing clonal populations, are well separated from the other cells within their ploidy fraction, both in terms of topology and branch lengths. This suggests the presence of additional clones or rare cell populations that were not previously reported.

**Figure 6.**
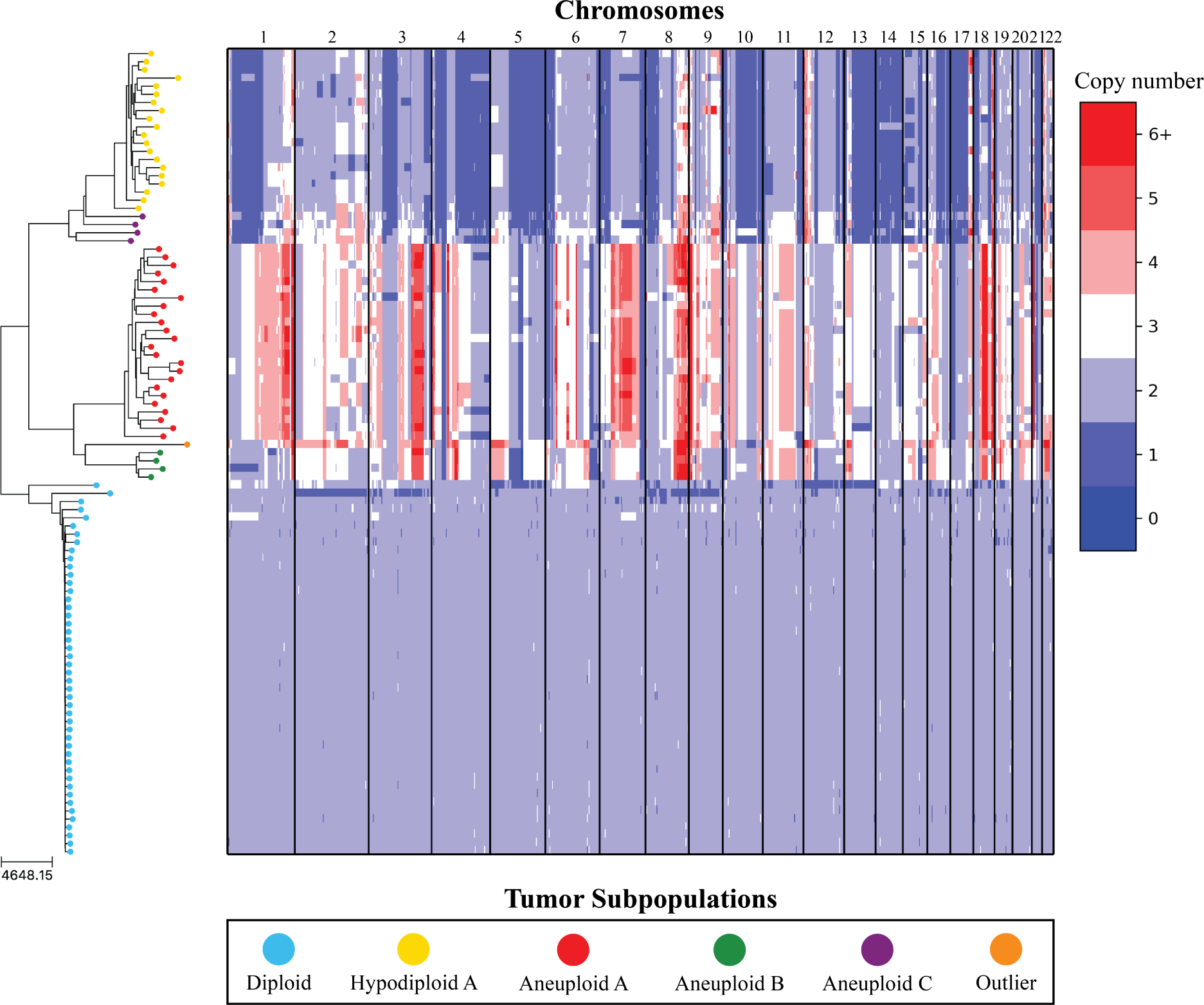
DICE-star tree of T10 aligned to a heatmap of whole-genome single-cell CNPs. Leaf nodes are shaded to match their corresponding clones derived by applying k-means clustering to the CNPs. The main clonal populations shown are defined by their ploidy: diploid (blue), hypodiploid (yellow), aneuploid A (red), and aneuploid B (green). Further increasing the number of clusters reveals an additional aneuploid clonal population (purple) from which the main hypodiploid population diverged, as well as a high-ploidy outlier cell (orange) placed in between aneuploid populations A and B. Both are distinguished by numerous chromosomal CNAs, suggesting that they represent additional rare or poorly sampled cell populations in the tumor.

To further investigate and validate these findings from the DICE-star trees, we identified clones independently by applying k-means clustering to the CNPs of each cell. We initially used a number of clusters equal to the major populations reported in Navin et al. (2011) (*k* = 4 for T10 and *k* = 3 for T16), and confirmed that this resulted in a near identical partitioning of the cell populations into distinct diploid, hypodiploid, and aneuploid fractions for T10, and distinct diploid, primary aneuploid, and metastatic aneuploid fractions for T16. These clusters also align with existing divergent clades in the DICE-star trees. Given the additional divergent clades/single nodes present in the DICE-star trees, we next investigated if using a different number of clusters would result in a more optimal clustering of the cells. We enumerated over all values 3*,…,*10 and found that using the number of clusters *k* = 6 and *k* = 5 resulted in the highest Silhouette Coefficient (Rousseeuw 1987) for T10 and T16, respectively. As expected, the updated cluster labels correspond to well-separated clades/branches of the DICE-star trees, strongly supporting the presence of previously unreported cell populations.

#### Analysis of T10

In patient T10, using *k* = 6 identifies two new small clusters in the DICE-star tree that were not identified by Navin et al. (2011). The first cluster is a 4-cell population with an average ploidy of 2.2 referred to as Aneuploid C (Figure 6). Given their ploidy, these are likely the same cells identified as pseudo-diploids by Navin et al. (2011) and found to be unrelated to one another in their analysis. However, as the DICE-star tree and the heatmap show (Figure 6), these 4 cells in Aneuploid C are likely closely related and form their own distinct population. The 4 cells undergo a number of shared large-scale aberrations, including amplifications on chromosomes 2, 11, 12, and 18, but also many of the same deletions as the hypodiploid population. In the DICE-star, the clade is positioned as a precursor clone from which the hypodiploid population originated. From the CNPs, we can observe that the hypodiploid population breaks away from Aneuploid C through an abundance of large-scale deletions, including back-mutations on virtually all amplified regions of Aneuploid C, which resulted in an average ploidy of 1.7. Given the evolutionary structure of the tree, and the fact that the average ploidy of Aneuploid C is over 2, this suggests that the hypodiploid population initially experienced copy number gains and losses in relatively equal abundance, followed by a reversion to the diploid state across multiple chromosomes and a single copy for others. The second cluster, a single outlier cell, is located in between Aneuploid populations A and B, made distinct by copy number gains on chromosomes 2p, 5p, 12, 15p, and 16p. The outlier cell has a higher ploidy than any other cluster at 3.3, compared to average ploidies of 3 and 2.8 for Aneuploid A and B. While the abnormal CNPs of the outlier cell may be the result of technical noise, this is unlikely since the cell passed the filtering protocol of SCOPE and harbors clear chromosomal CNAs.

#### Analysis of T16

In patient T16, using *k* = 5 retains the existing clonal populations, but identifies a new 3-cell cluster and an outlier cell on the DICE-star tree (Supplementary Figure S17) not previously identified by Navin et al. (2011). Both are well separated from other clones in the DICE-star tree. All cells in the 3-cell cluster, referred to as primary aneuploid B, are found in the primary site and have an average ploidy near 3. Interestingly, while one of the cells appears to have many unique copy number gains and losses, the other two appear to have a stable triploid state across the genome. This may indicate a stable yet poorly sampled cell population, but could also represent doublets or be an artifact of the CNA detection algorithm caused by normal cells with an abnormally high number of reads across the genome. The outlier cell, sampled from the metastatic site, has a high ploidy of 3.2 with erratic genome-wide profiles. While some regions of the genome are in extreme abundance (chromosomes 2, 7, and 8), some of which are shared with the other aneuploid cells, other regions uniquely have only a single copy (chromosomes 12 and 17).

## Discussion and Conclusion

In this work, we introduced two new methods, DICE-bar and DICE-star, for reconstructing tumor cell lineage trees from single-cell copy-number data. Both methods are based on novel, easy-to-compute distance measures, and drastically outperform the current state-of-the-art in terms of both accuracy and scalability. Using an extensive simulation study, we showed that DICE-bar matches or improves upon the accuracy of existing methods across all experimental conditions on noise-free data, and that DICE-star substantially improves upon all other methods, including DICE-bar, on nearly all datasets with noise/error levels similar to those observed in inferred CNPs on real sequence data. Remarkably, we also found that DICE-bar substantially outperforms far more sophisticated, MED-based methods across nearly all conditions and noise levels. Crucially, DICE-bar and DICE-star are not only more accurate than the other existing methods but also faster and more scalable.

The results of our simulation study also demonstrate the drastic effect that noise in CNPs has on the ability of the more sophisticated MED-based methods, based on nuanced models of copy number evolution, to effectively reconstruct the underlying phylogeny. At the same time, we find that DICE-star remains exceptionally tolerant to noise/error in the input CNPs, and dramatically outperforms those other methods for both low and high levels of noise. The lack of robustness to noise observed in MED-based methods is not entirely unexpected. Current limitations in scDNA-seq coverage necessitate the use of large bins to overcome poor resolution, and this can lead to single-cell CNPs that differ by only a small number of events. Consequently, even low levels of noise can disproportionately affect MED distance, particularly between cells that are closely related. This may explain why some MED-based methods, notably MEDICC2, appear to be more robust to noise when considering clone detection but show high sensitivity to noise when considering tree reconstruction accuracy. Still, the fact that DICE-bar, which is based on breakpoints and therefore similarly affected by noisy CNPs, outperforms existing MED-based methods on both noise-free and noisy datasets is surprising. Overall, one of the most important findings of this work is that relative simple distance-based methods, such as DICE-star or the method of Minussi et al. (Minussi et al. 2021), can produce more accurate tumor cell lineage trees on real scDNA-seq datasets than even the best existing MED-based or breakpoint based methods. Our analysis of two real breast cancer datasets highlights the potential impact of using more accurate tumor cell lineages reconstructed using DICE-star. On both datasets, analysis of the DICE-star cell lineage tree reveals possible rare cell populations that were previously unreported.

Our results suggest several important directions for future research on this problem. We found that the levels of noise typically observed in inferred CNPs on real single-cell datasets can greatly decrease the accuracy of all tumor cell lineage inference methods. Further improvements to methods for copy number estimation are therefore important for improving tumor cell lineage inference in practice. Our analysis suggests that DICE-star works very well overall on datasets across a wide range of noise levels. However, DICE-bar can deliver significant improvements over DICE-star when the data is relatively noise free. It may be possible to combine the distance functions of DICE-star and DICE-bar to design a new method that combines the strengths of both approaches. Likewise, it may be possible to adjust how breakpoints are computed to make DICE-bar more robust to noise. We also found that the MED-based method MEDICC2 performs very well when there are only a small number of bins or when the mutation rate is very low, as long as the data is noise free. It would thus be valuable to develop improved MED-based frameworks that are robust to noise. More robust MED-based methods should, in principle, be able to exceed the accuracies of much simpler methods like DICE-bar and DICE-star. Another promising direction is to apply a denoising and/or filtering step to the CNPs before computing distances, which was shown by sitka to be effective at handling noise under certain conditions.

Despite the accuracy and scalability of DICE-bar and DICE-star, their accuracy on very large datasets with thousands of cells may be limited by the local search heuristic used currently for reconstructing the final tree (balME implementation provided in FastME (Lefort et al. 2015)). The tree search may get stuck in local optima, limiting the accuracy of the methods. Improved tree search algorithms under minimum evolution, as well as strategies for escaping local optima, such as running replicates using resampled bins (analogous to resampling columns of a sequence alignment during phylogenetic bootstrapping), would therefore improve the accuracy of these methods on large datasets. Finally, some recent approaches have combined CNAs and single-nucleotide variants (SNVs) under a single model (Satas et al. 2020, Chen et al. 2022, Sollier et al. 2023, Zhang et al. 2023). SNVs, while more challenging to infer accurately from low-coverage single-cell sequencing data than CNAs (Rozhonova et al. 2022), could provide additional information and lead to more accurate tumor cell lineage trees.

## Data availability

The two breast cancer datasets of Navin et al. (2011) are available from the NCBI Sequence Read Archive under accession number SRA018951. All simulated datasets are freely available from Zenodo at https://doi.org/10.5281/zenodo.10108731. The software used for our analysis is freely available open-source from https://github.com/samsonweiner/DICE and https://compbio.engr.uconn.edu/software/dice/.

## Declaration of interests

The authors declare no competing interests.

## Acknowledgments

This work was supported in part by NSF award IIS 2212511.

## Notes

https://doi.org/10.5281/zenodo.10108731

https://github.com/samsonweiner/DICE

### Competing Interest Statement

The authors have declared no competing interest.

