## Supplementary text and supplementary figures for "DICE: Fast and Accurate Distance-Based Reconstruction of Single-Cell Copy Number Phylogenies"

#### S.1 Evaluation of alternative distance functions and phylogenetic reconstruction methods

**Selection of appropriate distance functions.** We explored the use of four simple and closely related distance functions: Euclidean (Eq. 1), Manhattan (Eq. 2), Root (Eq. 3), and Log (Eq. 4).

*Standard distances.* When applied to copy number profiles, we refer to these distances as 'standard' distances. Given two genomes $s$ and $t$ with $K$ chromosomes each, these standard distances are defined as follows:

$$d_{Euclidean}(s,t) = \sqrt{\sum_{k=1}^{K}\sum_{i=1}^{n_k}\left(c_{s,i}^{k} - c_{t,i}^{k}\right)^2}, \tag{1}$$

$$d_{Manhattan}(s,t) = \sum_{k=1}^{K}\sum_{i=1}^{n_k}\left|c_{s,i}^{k} - c_{t,i}^{k}\right|, \tag{2}$$

$$d_{Root}(s,t) = \sum_{k=1}^{K}\sum_{i=1}^{n_k}\sqrt{\left|c_{s,i}^{k} - c_{t,i}^{k}\right|}, \tag{3}$$

$$d_{Log}(s,t) = \sum_{k=1}^{K}\sum_{i=1}^{n_k}\log\left|c_{s,i}^{k} - c_{t,i}^{k}\right|. \tag{4}$$

*Breakpoint distances.* To address the limitations of standard distances, we proposed new distances functions that consider breakpoints rather than the copy numbers themselves. These breakpoint distances are defined analogously to their standard distance counterparts and their formal definitions appear below.

$$d'_{Euclidean}(s,t) = \sqrt{\sum_{k=1}^{K}\sum_{i=1}^{n_k-1}\left(b_{s,i}^{k} - b_{t,i}^{k}\right)^2}, \tag{5}$$

$$d'_{Manhattan}(s,t) = \sum_{k=1}^{K}\sum_{i=1}^{n_k-1}\left|b_{s,i}^{k} - b_{t,i}^{k}\right|, \tag{6}$$

$$d'_{Root}(s,t) = \sum_{k=1}^{K}\sum_{i=1}^{n_k-1}\sqrt{\left|b_{s,i}^{k} - b_{t,i}^{k}\right|}, \tag{7}$$

$$d'_{Log}(s,t) = \sum_{k=1}^{K}\sum_{i=1}^{n_k-1}\log\left|b_{s,i}^{k} - b_{t,i}^{k}\right|. \tag{8}$$

Of the *eight* distinct distance functions defined above, six are, to the best of our knowledge, novel (proposed and evaluated for the first time in this work). The other two, standard Euclidean and standard Manhattan, have previously been used to analyze real tumor samples (Navin et al. 2011, Minussi et al. 2021). The DICE software package implements all eight of these distance functions, and DICE-star and DICE-bar correspond to standard-Root (Eq. 3) and breakpoint-Root (Eq. 7), respectively.

**Distance-based phylogeny reconstruction.** We evaluated four possible distance-based phylogenetic reconstruction algorithms. These are Neighbor Joining (Saitou and Nei 1987), unweighted Neighbor Joining (Gascuel et al. 1997), balanced Minimum Evolution (Desper and Gascuel 2002), and ordinary least-squares Minimum Evolution (Rzhetsky and Nei 1993), and are referred to as NJ, uNJ, balME, and olsME, respectively. DICE uses the FastME software (Lefort et al. 2015), which implements all four of these methods, to compute the final cell lineage tree. FastME uses a local search heuristic for balME and olsME; an initial tree is built using an additive taxon procedure, and then an SPR tree search is used to find a topology minimizing the balME or olsME criteria.

**Evaluation of all 32 DICE variants.** We evaluated the 32 DICE variants (eight distance functions times four phylogeny reconstruction methods) using a subset of our simulated datasets. Specifically, we used our baseline datasets, simulated using CNAsim with the default parameter values and three different noise levels (no noise, low noise, and high noise) to assess the 32 DICE variants. We used these initial results to identify the best standard-distance method (DICE-star) and the best breakpoint-distance method (DICE-bar) for more thorough evaluation as described in the main manuscript.

Supplemental Figure S1 shows the results of this analysis. These results reveal interesting insights into the relative accuracies of the different DICE variants. First, we find that all Euclidean DICE variants, both standard and breakpoint, are among the worst performing methods at all noise levels. This includes the method of Navin et al. (2011), with corresponds to the standard-Euclidean-NJ variant of DICE. Second, we find that all non-Euclidean standard DICE variants show far greater accuracy than all other methods (including all breakpoint DICE variants) on noisy datasets. In contrast, on noise-free data, the non-Euclidean breakpoint DICE variants outperform all other methods. Third, among DICE variants, we find that the Root and Log distances show a slight advantage in performance over Manhattan distance, which includes the method of (Minussi et al. 2021), and that balME- and olsME-based variants perform better than the NJ- or uNJ-based ones across all noise levels. Based on these results, we selected the standard-root-balME variant of DICE as the best standard method DICE-star and the breakpoint-root-balME variant of DICE as the best breakpoint method DICE-bar.

We note that some other DICE variants, such as breakpoint-log-balME and standard-root-olsME, show nearly identical performance as DICE-bar and DICE-star, respectively. Among these equally strong variants, we chose root-balME as the basis for DICE-bar and DICE-star since the root distance function is slightly more interpretable than the log distance function, and since balanced minimum evolution (balME) has been observed to perform favorably compared to ordinary least-squares minimum evolution (olsME) in previous phylogenetic studies (Desper and Gascuel 2002; 2004).

#### S.2 Selection of noise parameters

The error rates used in CNAsim for generating noisy data were selected to reflect the performance of existing CNA detection methods reported in the benchmarking study of Mallory et al. (2020). This study computed the predicted copy number breakpoints of each method from sequencing data generated from simulated genomes where the ground truth copy number breakpoints are known. By comparing the genomic locations of the predicted and ground truth breakpoints, precision and recall values were derived for each of the evaluated methods. Across all experiments, the best performing methods in the study achieved precision and recall values in the ranges of $0.4-0.75$ and $0.5-0.8$, respectively.

For the selection of appropriate error rates in CNAsim, we chose values for the noise parameters $r_b$ and $r_j$ that result in precision and recall values within these ranges. When running CNAsim with a given combination of noise parameters, we are able to output the 'clean' CNPs in addition to the noisy CNPs. Thus, for a given combination of $r_b$ and $r_j$, we are able to compute a precision and recall values using the clean and noisy CNPs. In particular, given a clean CNP $C = (c_1, ..., c_n)$ and noisy CNP $C' = (c'_1, ..., c'_n)$, we apply the following procedure:

1. Compute the breakpoint profiles $B = (b_1, ..., b_{n-1})$ and $B' = (b'_1, ..., b'_{n-1})$.
2. Compute the sets of informative breakpoints $X = \{i : i \in 1...n-1 \text{ and } b_i \neq 0\}$ and $X' = \{i : i \in 1...n-1 \text{ and } b'_i \neq 0\}$.
3. Count the number of true positives (TP), false positives (FP), and false negatives (FN) as $|X \cap X'|$, $|X' - X|$, and $|X - X'|$, respectively, and compute precision and recall.

Table S1 details the precision and recall values of all noisy datasets generated with CNAsim.

We note that the relatively low precision and recall values for breakpoint detection on noisy CNPs (corresponding to those reported in Mallory et al. (2020)) are not caused by large-scale error in inferred/noisy copy numbers. In fact, the vast majority of copy numbers do not fluctuate and are left unchanged in the noisy CNPs. For example, in the boundary error model, since the number of contiguous segments (and therefore also breakpoints) is significantly less than the total number of bins, the boundary model alters a number of bins proportional to the number of segments. Likewise, the jitter error model works by redrawing each bin with copy number $c$ from a Normal distribution with a mean of $c$ and a standard deviation of $c * r_j$, where $r_j$ is the jitter error rate. This in effect means that the larger (resp. smaller) the copy number, the higher (resp. lower) the chances of observing jitter. For reference, an error rate of $r_j = 0.1$ will result in 9.55% of bins with copy number 3 to fluctuate, 1.24% of bins with copy number 2 to fluctuate, and a mere $5.73e^{-7}$% of bins with copy number 1 to fluctuate. On the default high noise setting (jitter error 0.1, boundary error 0.04), on average 1.05% of bins fluctuate compared to the noise-free profiles. On the default low noise setting, this value is 0.064%. These values match existing studies reporting high in-silico raw copy number recall (Funnell et al. 2022).

| Dataset | Boundary Error | Jitter Error | Precision | Recall |
|---|---|---|---|---|
| default low | 0.02 | 0.10 | 0.70969 | 0.76147 |
| default high | 0.04 | 0.10 | 0.65160 | 0.66836 |
| A1 | 0.02 | 0.05 | 0.80277 | 0.76819 |
| A2 | 0.04 | 0.05 | 0.73031 | 0.66376 |
| A3 | 0.06 | 0.05 | 0.67244 | 0.58627 |
| A4 | 0.08 | 0.05 | 0.63805 | 0.54182 |
| A5 | 0.10 | 0.05 | 0.61935 | 0.50606 |
| A6 | 0.15 | 0.05 | 0.55679 | 0.43148 |
| B1 | 0.02 | 0.15 | 0.57296 | 0.76853 |
| B2 | 0.04 | 0.15 | 0.50085 | 0.66409 |
| B3 | 0.06 | 0.15 | 0.45283 | 0.59533 |
| B4 | 0.08 | 0.15 | 0.4338 | 0.55597 |
| B5 | 0.10 | 0.15 | 0.40798 | 0.51723 |
| B6 | 0.15 | 0.15 | 0.35045 | 0.45255 |

**Table S1. Precision and recall values for various combinations of noise parameters.** The reported precision and recall values are estimated by comparing ground truth CNPs with the generated, noisy CNPs as described in Supplementary Section S.2. Datasets A1-A6 correspond to those depicted in Figure **??** Part (a), while datasets B1-B6 correspond to those depicted in Figure **??** Part (b). All reported values are averaged over 20 datasets.

#### S.3 Key differences between the three simulation frameworks

We simulated datasets using CNAsim (Weiner and Bansal 2023) and the simulation frameworks of two previous methods MEDICC2 (Kaufmann et al. 2022) and cnp2cnp (Cordonnier and Lafond 2020). All three simulation frameworks take the same general approach: A ground truth tree topology is generated, CNAs are accumulated along each edge, and observed CNPs are derived from the genomes of the leaf nodes. However, there are several important differences between the two previous simulation frameworks and CNAsim. Some of the key differences include the following.

1. Number of bins: The simulation study of MEDICC2 used at most 10 bins per chromosome with $2 \times 22$ chromosomes, for a total of 440 bins, and the simulation study of cnp2cnp used at most 250 bins from a single chromosome. In contrast, datasets generated by CNAsim have a more realistic number of bins by default which, at 1 Mbp bin lengths and the $2 \times 22$ autosome lengths of hg38, comes out to roughly 5760 bins.
2. Resolution of genome: Both MEDICC2 and cnp2cnp simulation frameworks simulate events over the bins themselves, meaning CNAs always start and end exactly at the boundaries between bins. For comparison, CNAsim simulates events at the region level, which are by default 1000x smaller than the bins. This results in more realistic CNAs when using CNAsim since real CNAs very rarely start or end at bin boundaries or may be contained within a single bin entirely.
3. Magnitudes of copy number gains: Both of the older simulation frameworks model deletions and duplications, but not copy number amplifications of larger magnitudes. CNAsim models the magnitude of a copy number amplification with a geometric distribution such that the higher the magnitude, the lower the probability. This results in duplications still being the most common, but higher magnitude events are possible.
4. Simulation of the cell-lineage tree: Both MEDICC2 and cnp2cnp simulation frameworks construct the ground truth topology by randomly merging cells, compared to the coalescent model used in CNAsim.
5. Zero copy number: While CNAsim allows the deletion of the last copy of a region, cnp2cnp considers various rates of this occurring, and MEDICC2 disallows it completely.
6. Error in copy number profiles: The simulation framework of cnp2cnp has a simple noise-model similar to the jitter model of CNAsim. The simulation framework of MEDICC2 can only generate noise-free CNPs and does not explore noisy copy number profiles. CNAsim has the most complex noise model and includes both jitter error and boundary error.
7. Event types: The simulation framework of cnp2cnp only uses a single chromosome, and is limited to segmental duplications and deletions. The simulation framework of MEDICC2 models a variety of events including segmental duplications and deletions, WGD, and whole-chromosomal duplications and deletions. Additionally, MEDICC2 models several events that are not included in CNAsim, namely balanced and unbalanced translocations, insertions, and inversions.

#### S.4 Specific commands used for running existing methods

The specific commands and scripts used for running MEDICC2 (Kaufmann et al. 2022), MEDALT (Wang et al. 2021), cnp2cnp (Cordonnier and Lafond 2020), sitka (Salehi et al. 2023), and Lazac (Schmidt et al. 2023) in our exprimental analysis are available in the DICE GitHub repository (https://github.com/samsonweiner/DICE). These methods were all run in accordance to their respective software manuals or readme files using suggested or default options. For WCND (Zeira and Raphael 2020), there does not exist a fully operable software package or user manual and we therefore describe how we used WCND below.

The WCND code repository on GitHub provides only high level functions and does not include examples or a guide to defining weight functions. Because of this, we used the "semi_directed_cnd" function to compute unweighted MED distances over all chromosomal CNPs of each pair of cells, set the total distance between a pair of cells as the sum of the chromosomal unweighted MED distances, and reconstructed the tree from the pairwise distances using neighbor joining.

Finally, we note that MEDICC2 also provides options for reconstructing ancestors on the inferred phylogeny. To enable a fair comparison of running times, we enabled the *–topology-only* flag of MEDICC2 so as to output the tree topology only (skips reconstructing ancestors), and also enabled the *–no-plot* flag to skip generating visual plots.

### Supplementary Figures

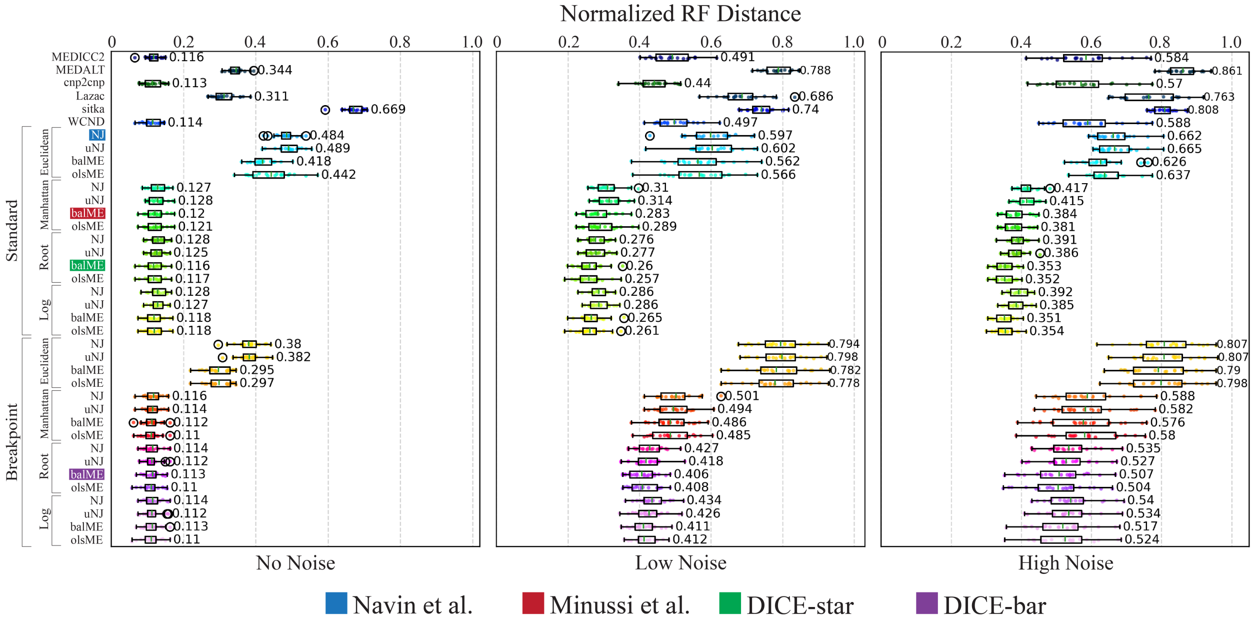


**Figure S1. Reconstruction accuracies of all 32 DICE variants and existing methods.** Cell lineage tree reconstruction accuracies are shown for all 32 DICE variants and existing methods on simulated datasets with 250 cells and varying levels of noise. The datasets correspond to those appearing in Figure 2 of the main text. Results for datasets with no noise, low noise, and high noise appear in the left, middle, and right panels, respectively. The methods of Navin et al. and Minussi et al., which correspond to two of the DICE variants, are highlighted in blue and red, respectively. All results are averaged over 20 simulated datasets.

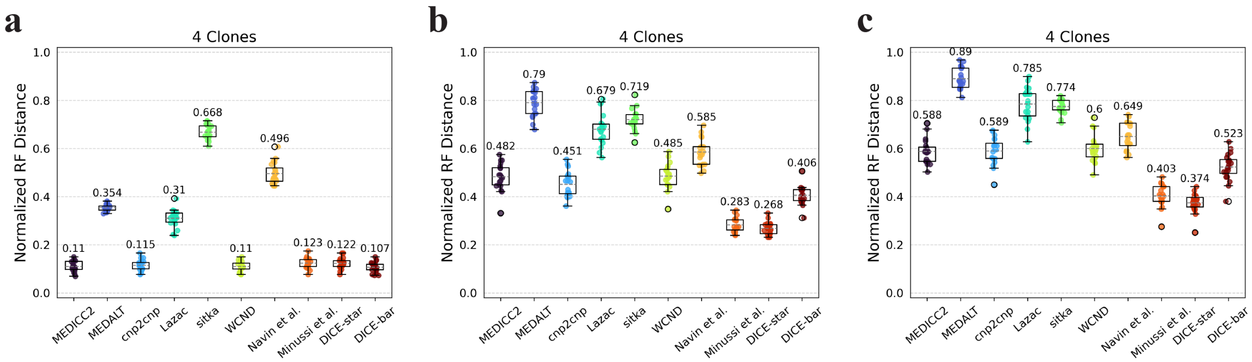


**Figure S2. Cell lineage tree reconstruction accuracy on four-clone trees.** Cell lineage tree reconstruction accuracies are shown for the datasets used in Figure 3 of the main text. Observe that performance across all methods is largely unaffected by the presence of clonal structures, with DICE-bar continuing to outperform all other methods on noise-free datasets and DICE-star continuing to substantially outperform all other methods on the low- and high-noise datasets. Scores are averaged over 20 datasets, each with exactly four clones.

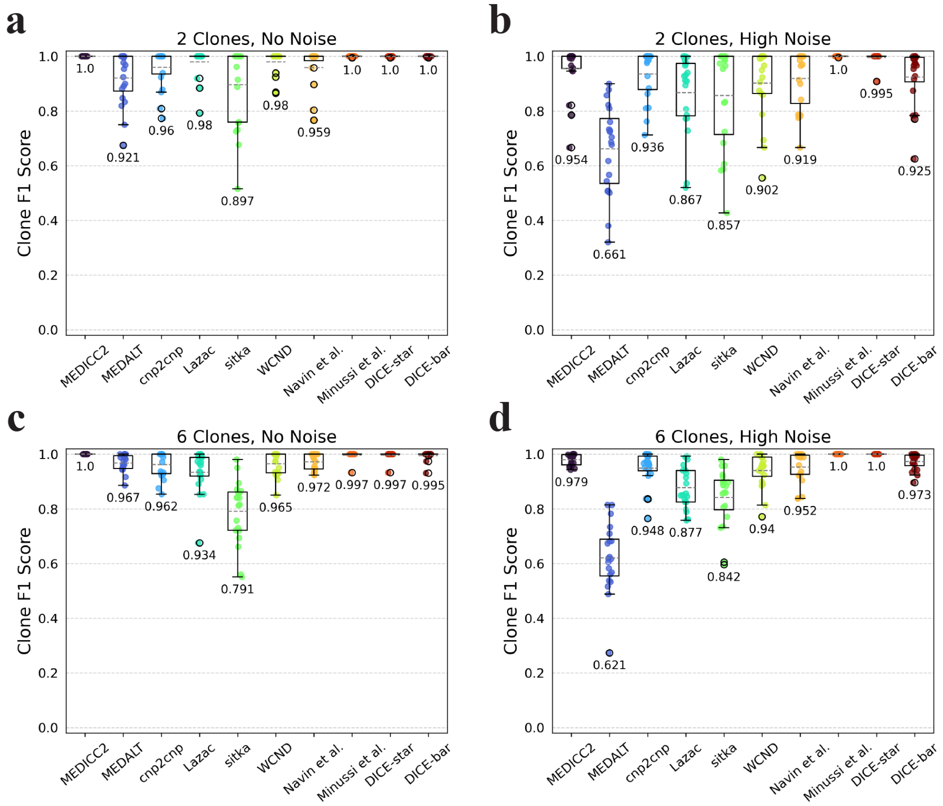


**Figure S3. Clone detection accuracies on datasets with varying numbers of clones**. F1 scores are shown for the different methods on simulated datasets with 250 cells and either no noise (left column) or high noise (right column). Scores are averaged over 20 datasets, each with exactly two (top row) or exactly six (bottom row) clones. Observe that these results are consistent with those shown in Figure 3 in the main text which shows results for exactly four clones.

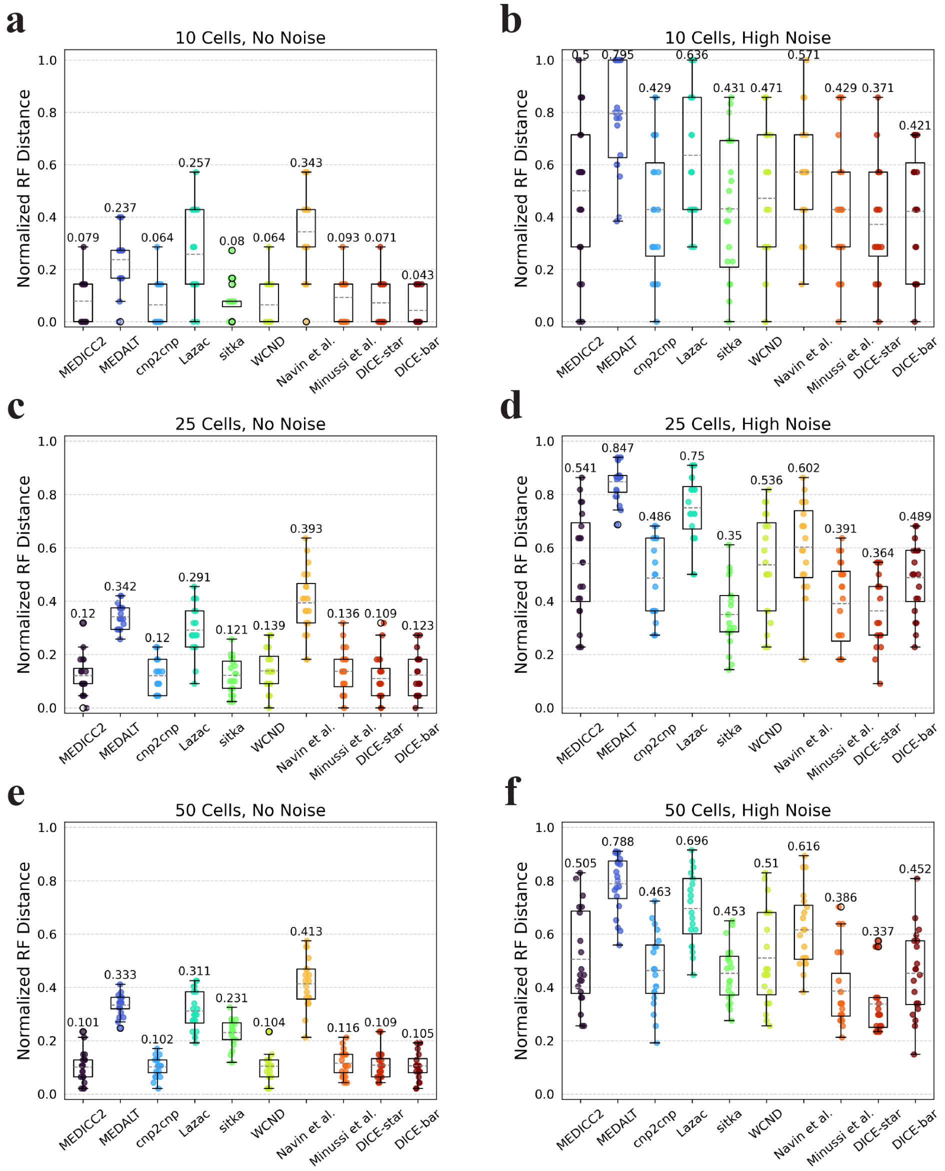


**Figure S4. Reconstruction accuracy for smaller numbers of cells.** Cell lineage tree reconstruction accuracies are shown for different methods on simulated datasets with varying numbers of cells (rows; 10, 25, and 50 cells) and at two different noise levels (columns; no-noise and high-noise). Lower normalized RF distances imply greater tree reconstruction accuracy. All reported results are averaged over 20 datasets.

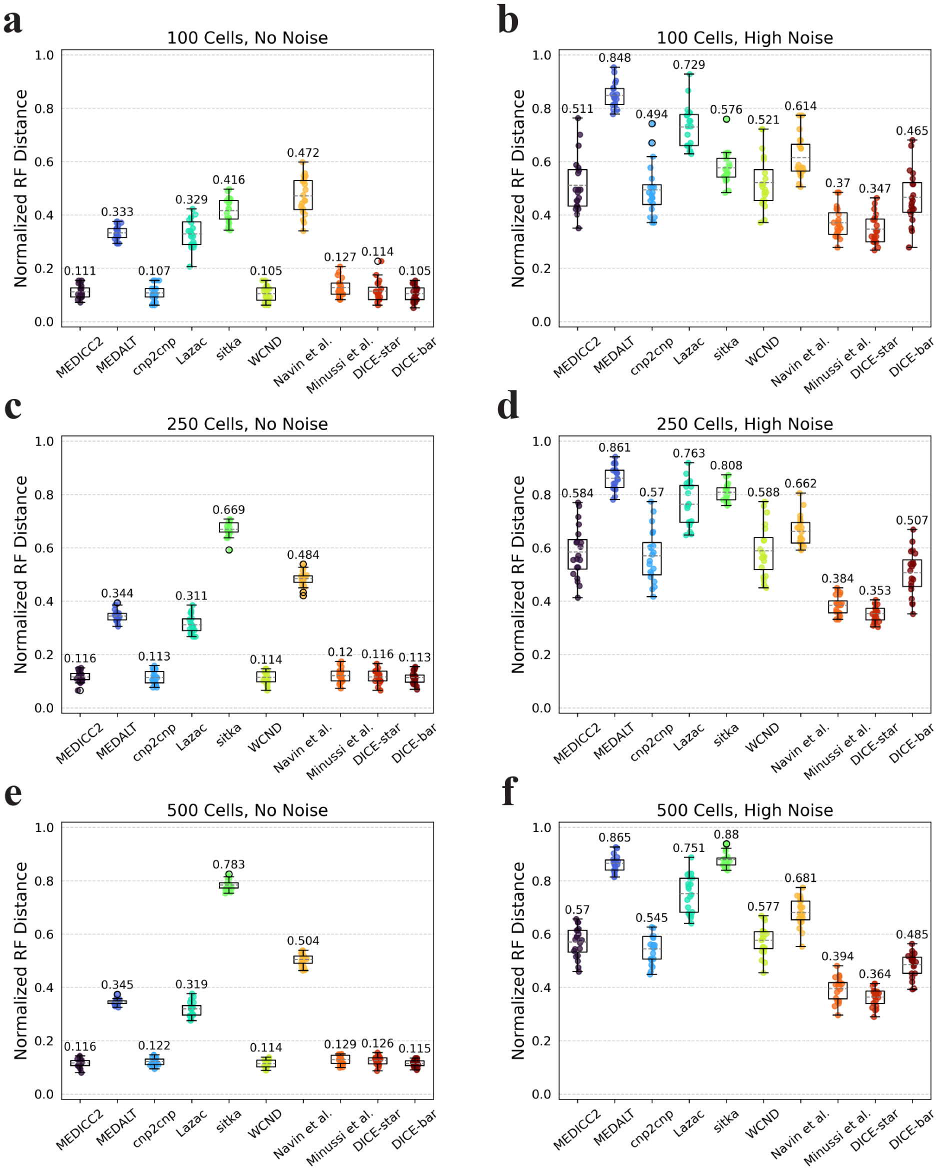


**Figure S5. Reconstruction accuracy for larger numbers of cells.** Cell lineage tree reconstruction accuracies are shown for different methods on simulated datasets with varying numbers of cells (rows; 100, 250, and 500 cells) and at two different noise levels (columns; no-noise and high-noise). Lower normalized RF distances imply greater tree reconstruction accuracy. All reported results are averaged over 20 datasets.

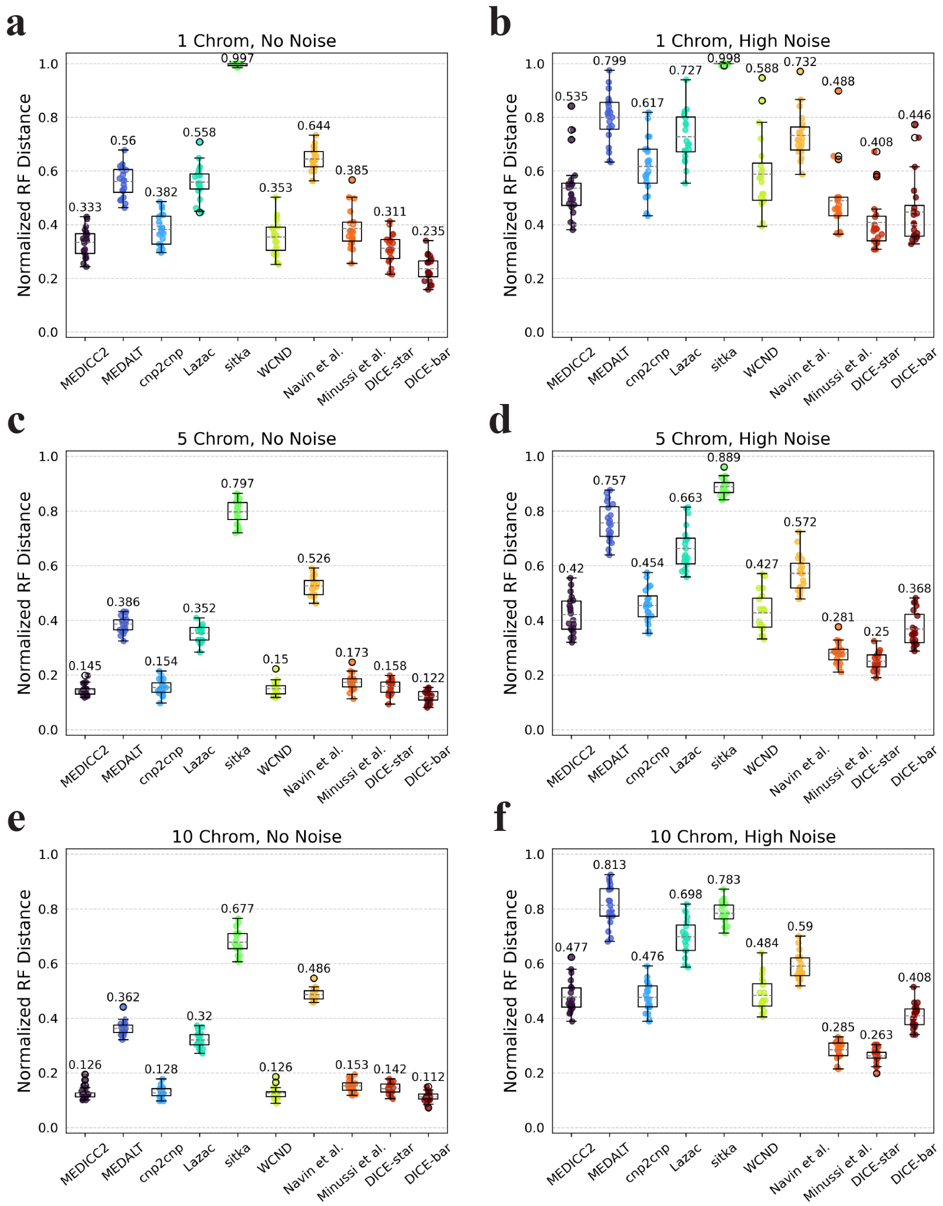

**Figure S6. Impact of number of chromosomes on reconstruction accuracy.** Cell lineage tree reconstruction accuracies are shown for different methods on simulated datasets with varying numbers of chromosomes (rows; 1, 5, and 10 chromosomes) and at two different noise levels (columns; no-noise and high-noise). Lower normalized RF distances imply greater tree reconstruction accuracy. All reported results are averaged over 20 datasets.

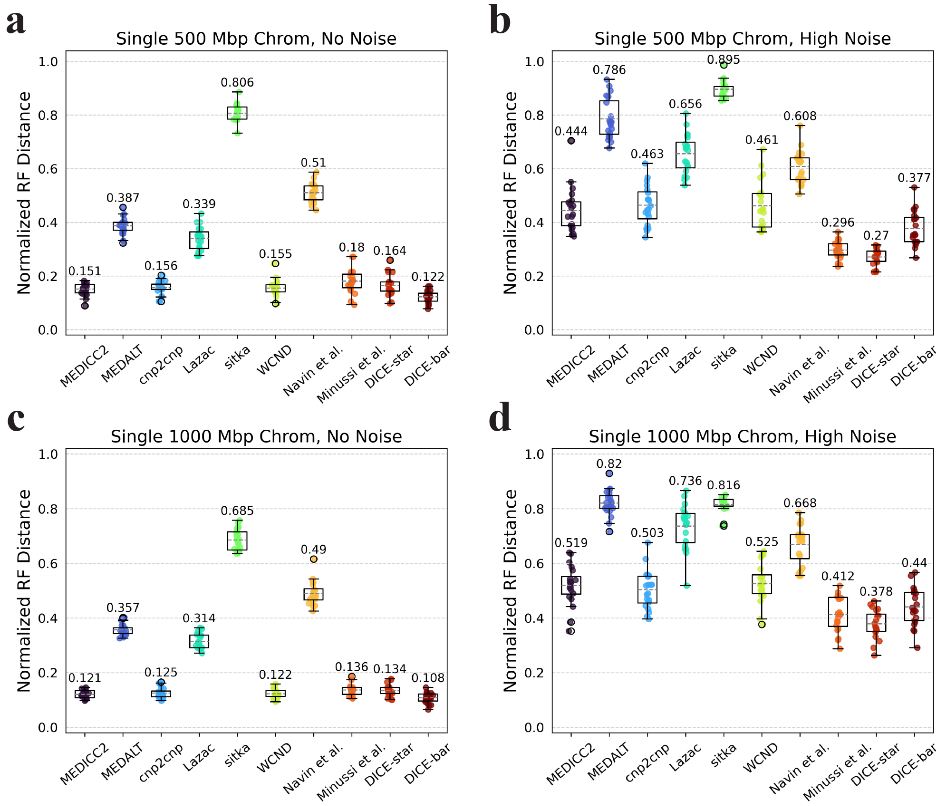


**Figure S7. Reconstruction accuracy on single chromosomes of variable length.** Cell lineage tree reconstruction accuracies are shown for different methods on simulated datasets with chromosomes of variable length (rows; 500 Mbp and 1000 Mbp chromosome lengths) and at two different noise levels (columns; no-noise and high-noise). Lower normalized RF distances imply greater tree reconstruction accuracy. All reported results are averaged over 20 datasets.

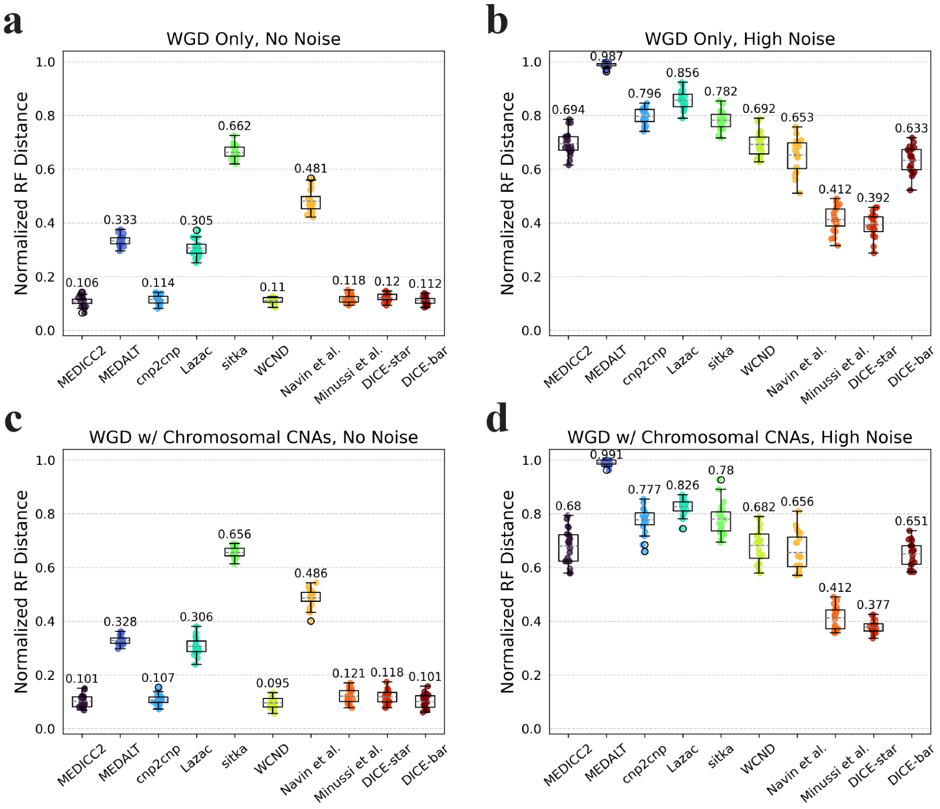


**Figure S8. Reconstruction accuracy in the presence of WGD.** Cell lineage tree reconstruction accuracies are shown for different methods on simulated datasets with WGD (rows) and at two different noise levels (columns; no-noise and high-noise). WGD was evaluated both individually (top row) and in the presence of chromosomal CNAs (bottom row). For the latter, a mean of 2 chromosomal events occur on the edges into and out of the root node, and into 4 clonal clades existing in the phylogeny. Lower normalized RF distances imply greater tree reconstruction accuracy. On noise-free data, performance slightly improves when chromosomal CNAs are included, but is otherwise unaffected by WGD. On noisy data, Minussi et al. and DICE-star perform slightly worse in the presence of WGD, while all other methods suffer heavily from the increase in ploidy due to WGD. All reported results are averaged over 20 datasets.

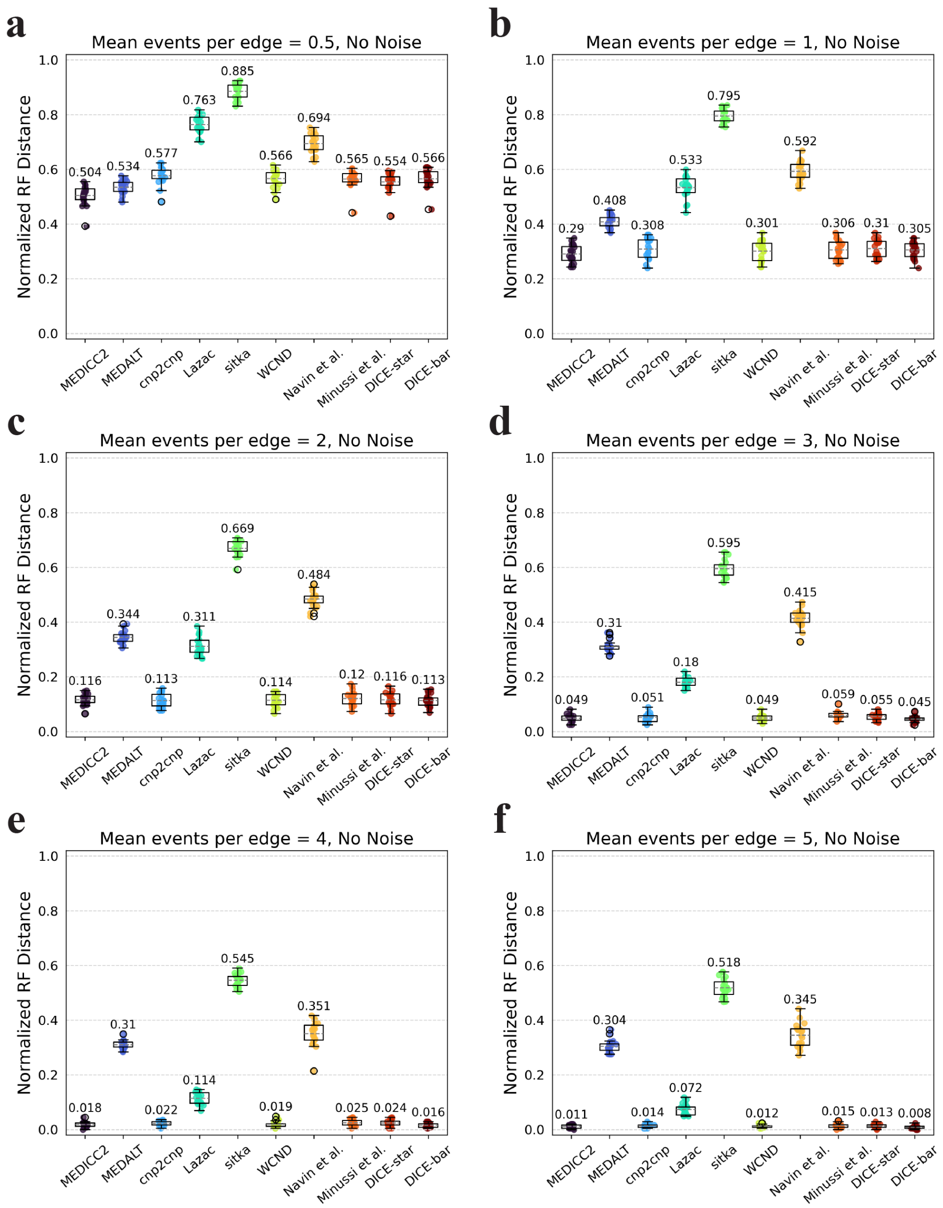


**Figure S9. Impact of events per edge on reconstruction accuracy using noise-free data.** Cell lineage tree reconstruction accuracies are shown for different methods on simulated noise-free datasets with increasing numbers of focal CNAs per edge (0.5, 1, 2, 3, 4, 5 events per edge). Lower normalized RF distances imply greater tree reconstruction accuracy. For every one additional event per edge, the normalised RF distance roughly halves for all methods except MEDALT, sitka, and Navin et al. All reported results are averaged over 20 datasets.

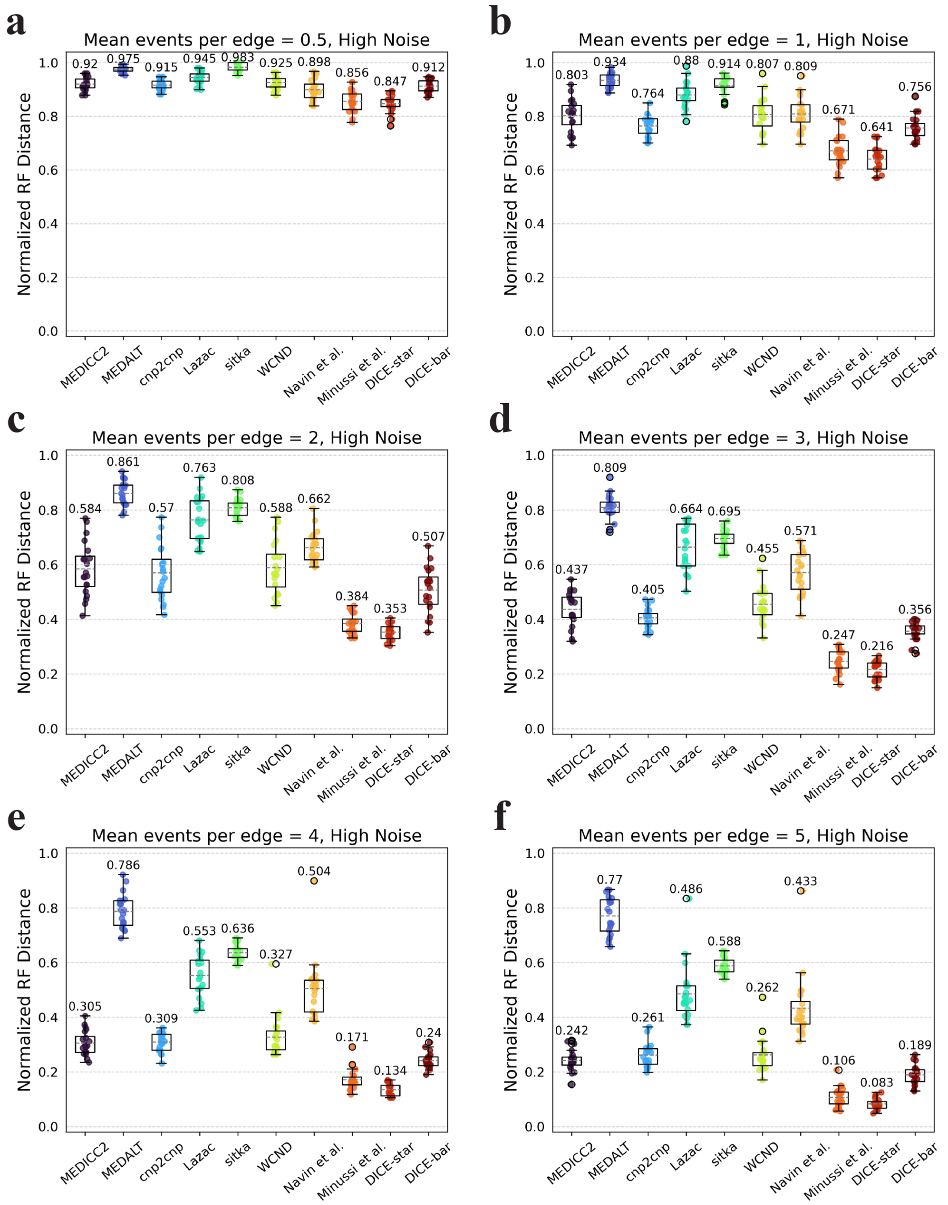


**Figure S10. Impact of events per edge on reconstruction accuracy using noisy data.** Cell lineage tree reconstruction accuracies are shown for different methods on simulated noisy datasets with increasing numbers of focal CNAs per edge (0.5, 1, 2, 3, 4, 5 events per edge). Lower normalized RF distances imply greater tree reconstruction accuracy. These datasets correspond to those depicted in Figure S9 using a high noise rate. Increasing the mutation rate substantially improves the performance of all methods except for MEDALT and sitka, which only improve slightly. Observe that the effect of noise is amplified under very low mutations rates (parts a and b), as the CNA-to-noise ratio is severely imbalanced. All reported results are averaged over 20 datasets.

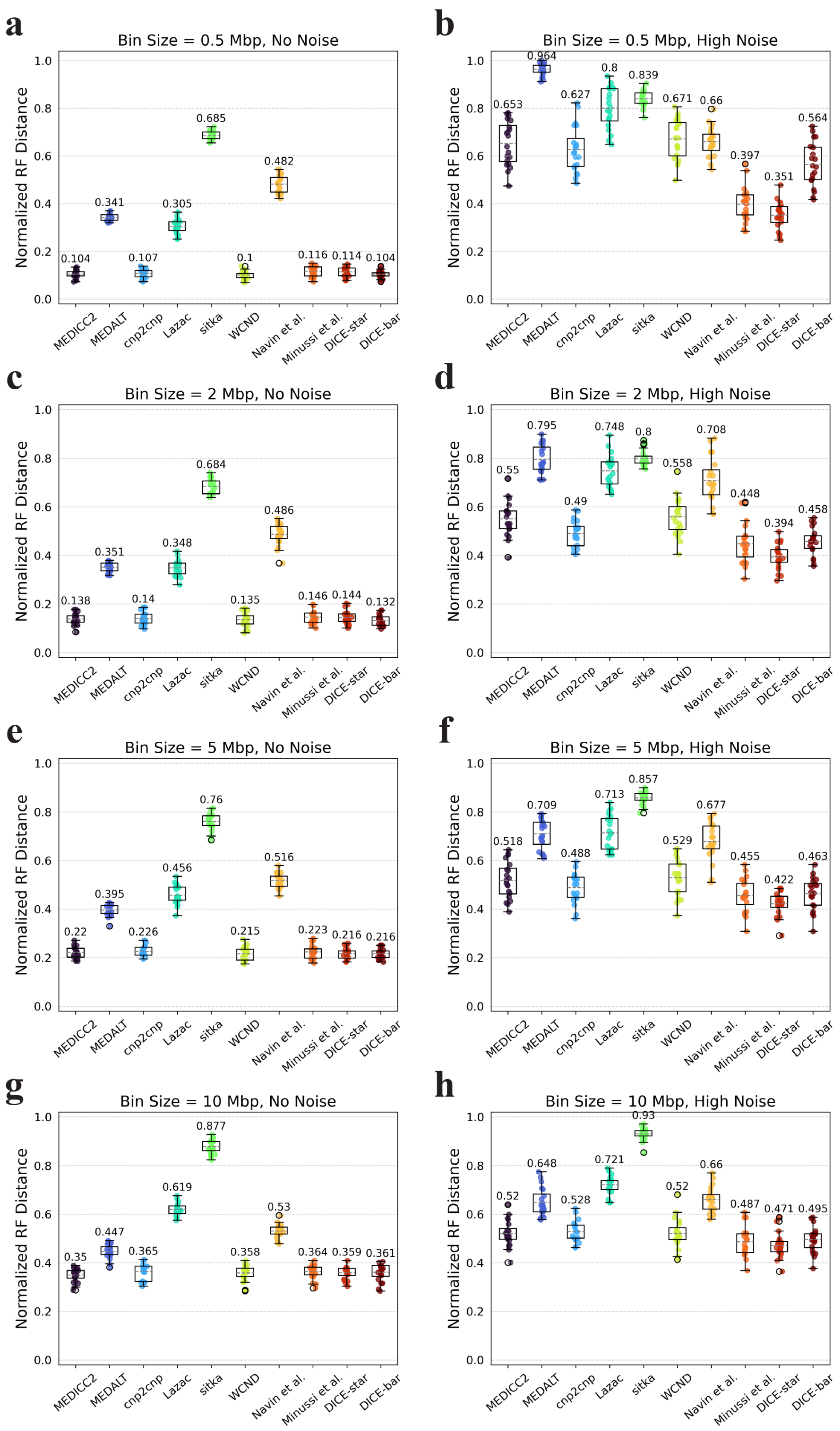


**Figure S11. Impact of bin size on reconstruction accuracy.** Cell lineage reconstruction accuracies are shown for different methods on simulated datasets with increasing bin size (rows; 0.5 Mbp, 2 Mbp, 5 Mbp, and 10 Mbp bin sizes) and at two different noise levels (columns; no-noise and high-noise). Lower normalized RF distances imply greater tree reconstruction accuracy. On noise-free datasets, across all methods, performance steadily decreases with an increase in bin size. This trend is, surprisingly, large reversed on noisy datasets, with only DICE-star and Minussi et al. clearly benefiting from smaller bin sizes. All reported results are averaged over 20 datasets.

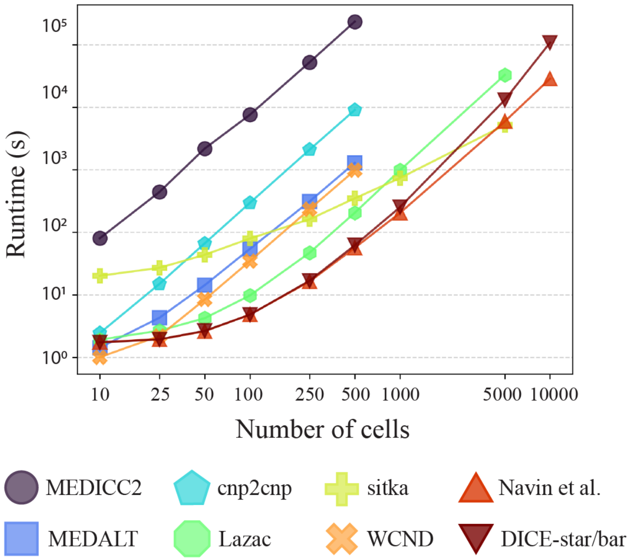


**Figure S12. Running time and scalability on noisy data.** Running times are shown for MEDICC2, MEDALT, cnp2cnp, Lazac, sitka, WCND, Navin et al., Minussi et al., DICE-bar, and DICE-star on high-noise datasets with varying numbers of cells. Running times for DICE-bar, DICE-star, and the method of Minussi et al. (2021) are identical and are shown together. The *x*-axis is the log-scaled number of cells in the dataset, and the *y*-axis is the log-scaled runtime in seconds. All reported times are averaged over 20 runs executed using a single core of an Intel Xeon 2.1 GHz processor.

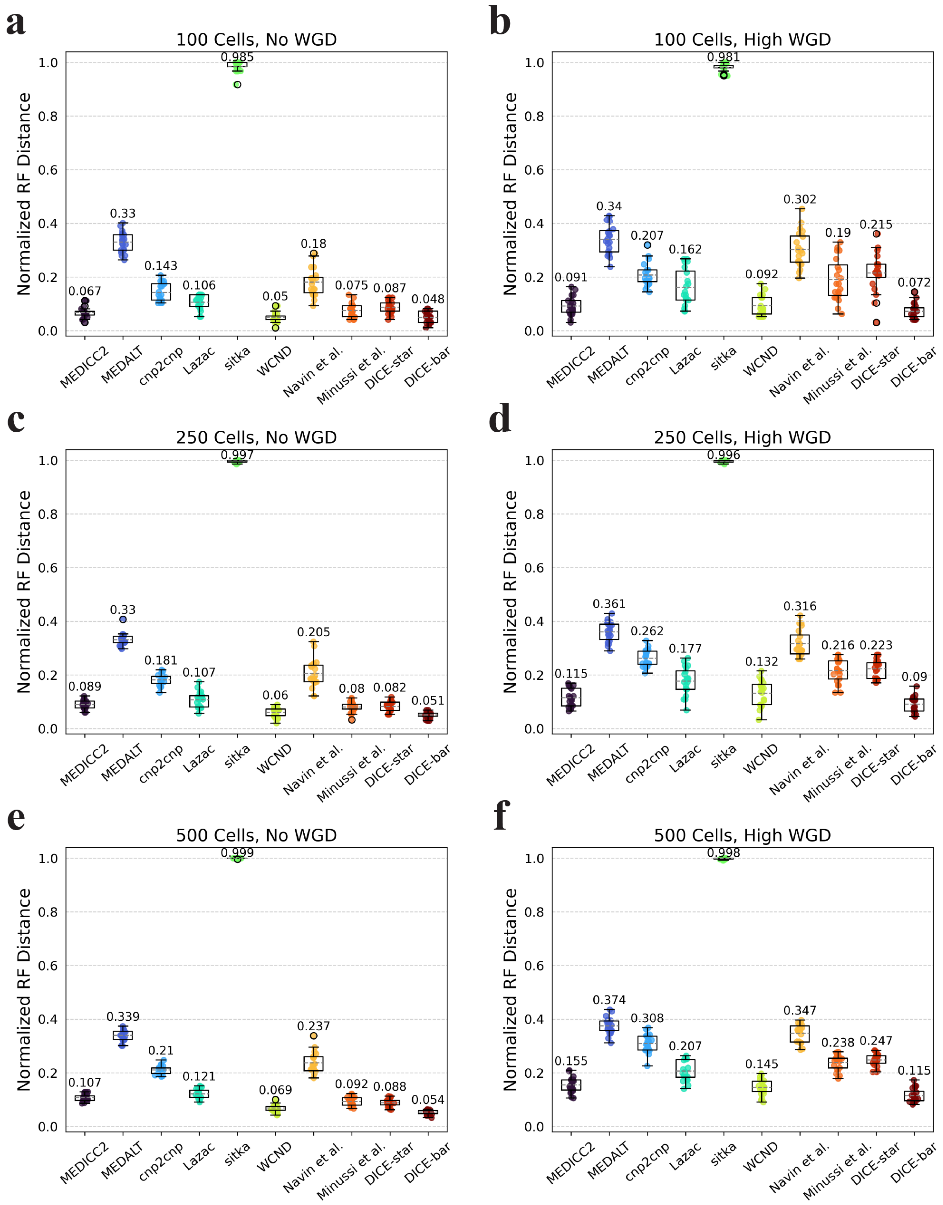


**Figure S13. Results using MEDICC2's simulation framework with default parameters.** Cell lineage reconstruction accuracies are shown for different methods on datasets simulated using the simulation routine of MEDICC2 with default parameters and no noise. Left column: (**a, c, e**) results on datasets with WGD rate of 0 and an increasing number of cells. Right column: (**b, d, f**) Results on datasets with high WGD rate and increasing number of cells. Simulation parameter values were selected as those described in (Kaufmann et al. 2022). The results show similar trends as those on noise-free simulated datasets of CNAsim, with DICE-bar significantly outperforming all other methods.

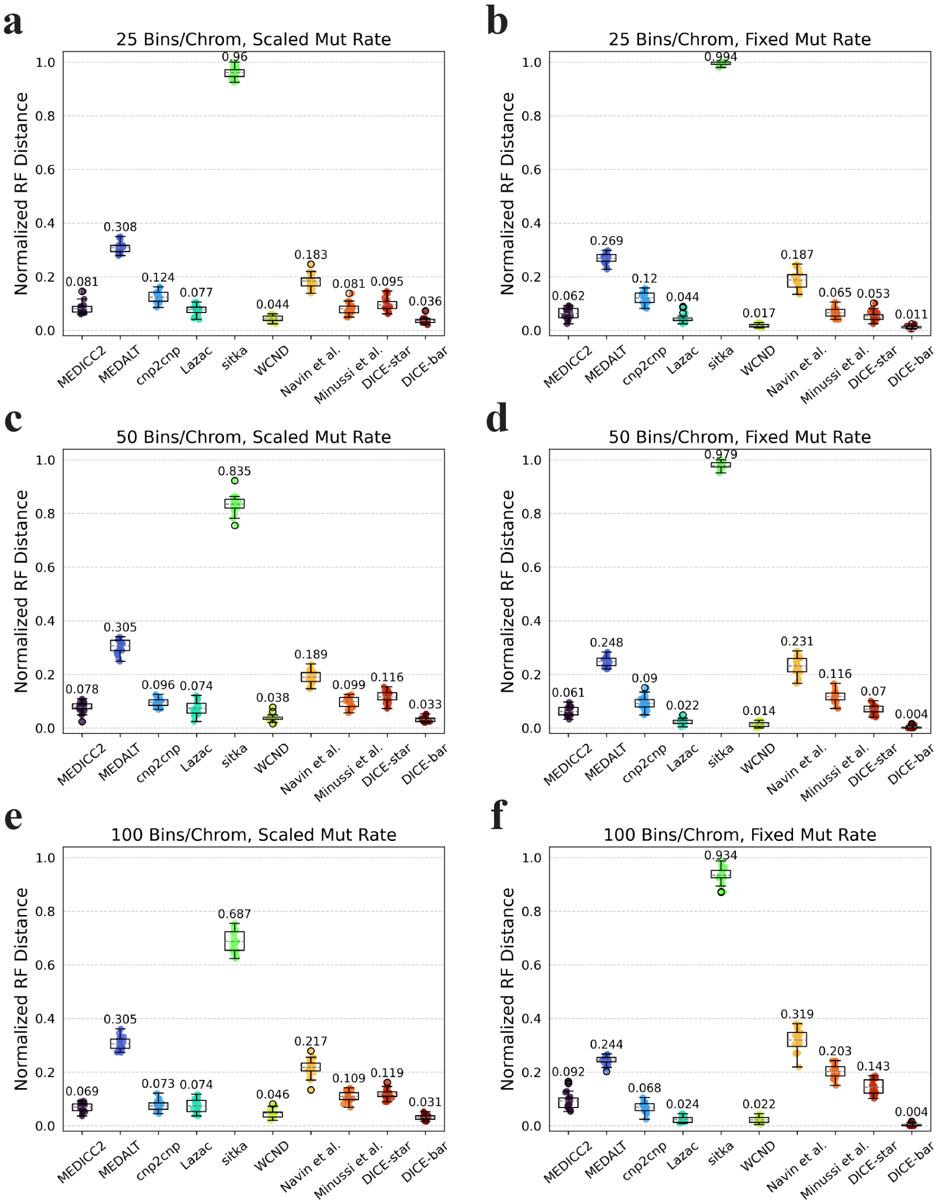


**Figure S14. Results using MEDICC2's simulation framework with increasing number of bins per chromosome.** Cell lineage reconstruction accuracies are shown for different methods on datasets simulated using the simulation routine of MEDICC2 with default parameters, no noise, and increasing numbers of bins (rows; 25, 50, and 100 bins per chromosome). Left column: (**a, c, e**) Performance on simulated datasets of increasing number of bins per chromosome with a scaled mutation rate such that the number of events per edge is consistent across all settings. Right column: (**b, d, f**) Performance on simulated datasets of increasing number of bins per chromosome with a fixed mutation rate of 0.05. Note that the number of events per edge in the MEDICC2 simulation routine scales with total genome size, so datasets of varying genome size cannot be consistently compared when using a fixed mutation rate. Observe that DICE-bar drastically outperforms all other methods on these datasets. All reported results are averaged over 20 datasets.

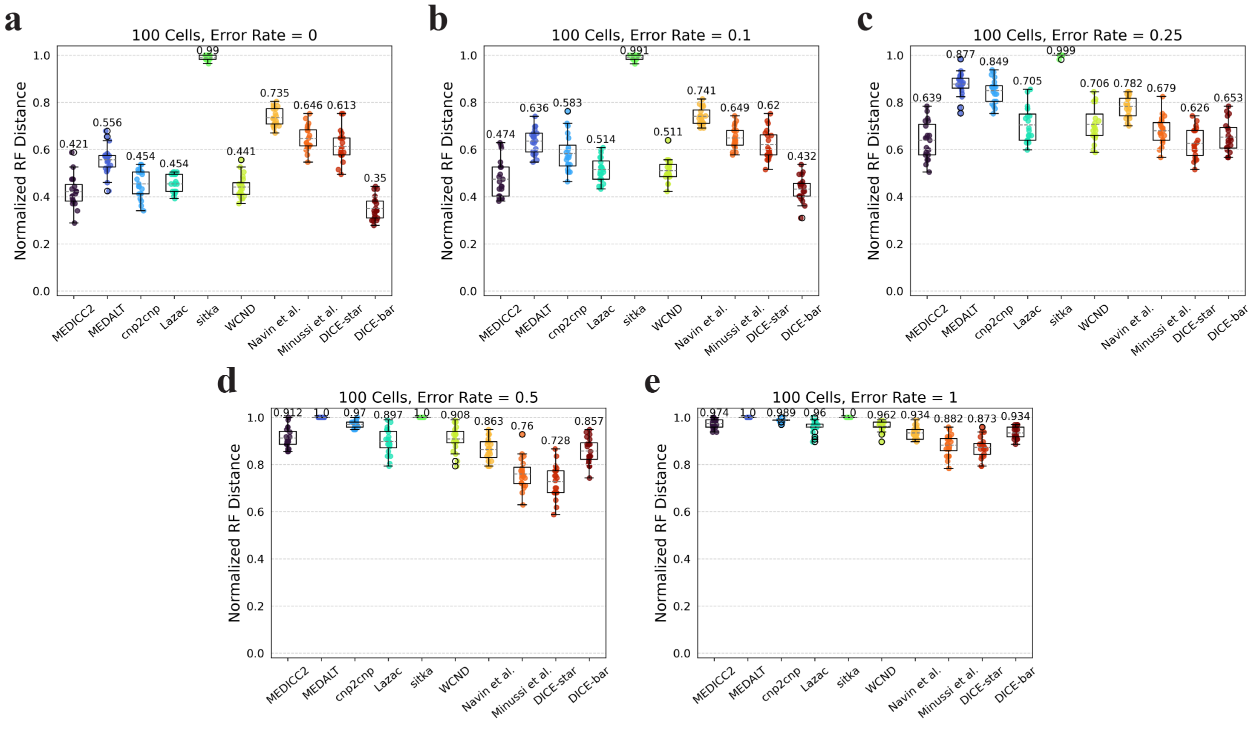


**Figure S15. Results using cnp2cnp's simulation framework with default parameters.** Cell lineage reconstruction accuracies are shown for different methods on 100-cell datasets simulated using the simulation routine of cnp2cnp with default parameters and increasing levels of noise (error rates). Observe that the performance of all methods is significantly worse compared to performance on the simulated datasets of CNAsim, likely due to the very low default number of bins (set to 100 on a single chromosome). DICE-bar outperforms all other methods for the two lowest noise levels, while DICE-star outperforms all other methods for all remaining (higher) noise levels. All results are averaged over 20 datasets.

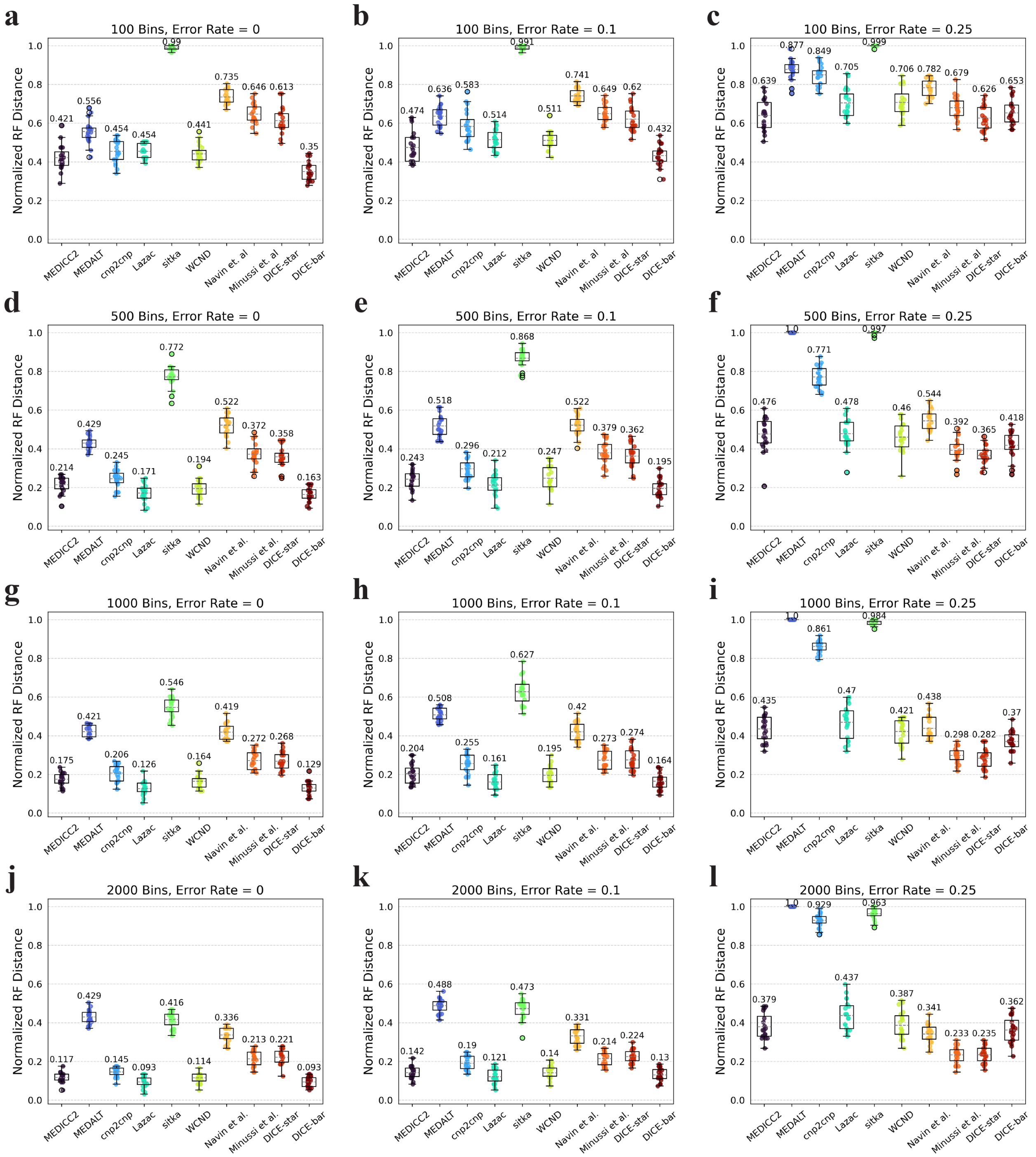


**Figure S16. Results using cnp2cnp's simulation framework with increasing number of bins.** Cell lineage reconstruction accuracies are shown for different methods on 100-cell datasets simulated using the simulation routine of cnp2cnp with increasing numbers of bins (rows; 10, 500, 1000, and 2000 bins) and at three different noise levels (columns; error rates 0, 0.1, and 0.25). As the number of bins increases, for most methods, the overall performance of all methods steadily increases. The relative performance of methods remains unchanged, with either DICE-bar or DICE-star matching or outperforming the other methods depending on the level of noise in the dataset. All results are averaged over 20 datasets.

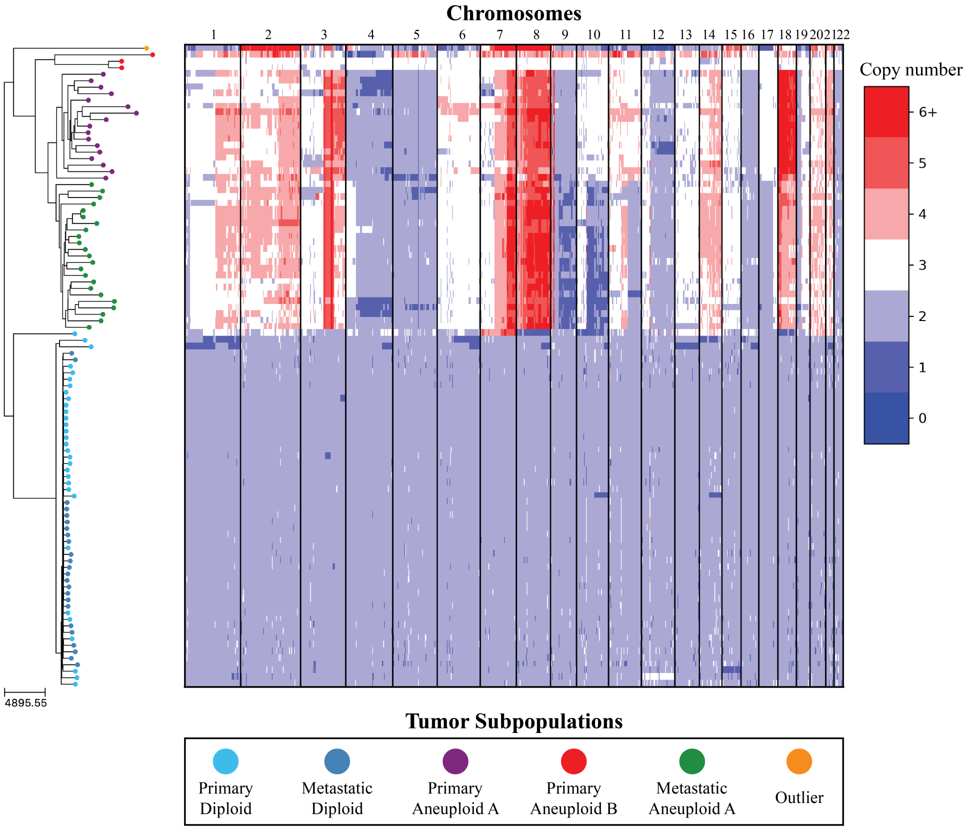


**Figure S17. DICE-star tree of T16 aligned to a heatmap of whole-genome copy number profiles.** Leaf nodes are shaded to match their corresponding clones derived from applying k-means clustering to the copy number profiles. The main clonal populations displayed are defined by their ploidy: diploid (blue), primary aneuploid A (purple), and metastatic aneuploid (green). Further increasing the number of clusters reveals an additional small aneuploid population from the primary site (red) that is significantly well-separated from the other clones, and an outlier cell sampled from the metastatic site whose copy number profiles contain a multitude of unique chromosomal CNAs (orange). Both appear to have diverged from the previously reported populations far earlier than with each other, and may represent undersampled rare populations.
